# Plant defensin MtDef4-derived antifungal peptide with multiple modes of action and potential as a bioinspired fungicide

**DOI:** 10.1101/2022.10.02.510465

**Authors:** Meenakshi Tetorya, Hui Li, Arnaud Thierry Djami-Tchatchou, Garry W. Buchko, Kirk J. Czymmek, Dilip M. Shah

## Abstract

Chemical fungicides have been instrumental in protecting crops from fungal diseases. However, mounting fungal resistance to many of the single-site chemical fungicides calls for the development of new antifungal agents with novel modes of action (MoA). The sequence-divergent cysteine-rich antifungal defensins with multi-site MoA are promising starting templates for design of novel peptide-based fungicides. Here, we experimentally tested such a set of 17-amino acid peptides containing the γ-core motif of the antifungal plant defensin MtDef4. These designed peptides exhibited antifungal properties different from those of MtDef4. Focused analysis of a lead peptide, GMA4CG_V6, showed it was a random coil in solution with little or no secondary structure elements. Additionally, it exhibited potent cation-tolerant antifungal activity against the plant fungal pathogen *Botrytis cinerea*, causal agent of gray mold disease in fruits and vegetables. Its multi-site MoA involved localization predominantly to the plasma membrane, permeabilization of the plasma membrane, rapid internalization into the vacuole and cytoplasm, and affinity for bioactive phosphoinositides phosphatidylinositol 3-phosphate (PI3P), PI4P, and PI5P. The sequence motif RRRW was identified as a major determinant of the antifungal activity of this peptide. While topical spray-application of GMA4CG_V6 on *Nicotiana benthamiana* and tomato plants provided preventative and curative suppression of gray mold disease symptoms, the peptide was not internalized into plant cells. Our findings open the possibility that truncated and modified defensin-derived peptides containing the γ-core sequence could serve as promising candidates for further development as bioinspired fungicides.

## INTRODUCTION

Crop plants are under constant threat from fungal and oomycete pathogens that cause significant pre-and post-harvest economic losses worldwide and challenge food security globally (Fones *et al*., 2020). This threat is increasing in severity and scale (Fisher *et al*., 2012, Savary *et al*., 2019). It is estimated that fungal and oomycete pathogens cause up to 10-23% yield losses in calorie crops (Savary et al., 2019) and postharvest losses of 10% (Fisher *et al*., 2018). Modern intensive agriculture depends heavily on chemical fungicides for management of fungal and oomycete diseases in commodity crops as well as high value vegetable and fruit crops (Oliver & Hewitt, 2014). Most chemical fungicides in use today have a single biochemical target and thus are prone to resistance development by diverse and rapidly evolving fungal and oomycete strains (Hawkins and Fraiije, 2018). Facing fungal resistance and unwanted side effects, some fungicides are losing their effectiveness in modern intensive monoculture cropping systems. The structurally diverse plant antifungal peptides with potent antifungal activity could potentially serve as structural templates for the design of a new generation of safe and sustainable fungicides (Lobo & Boto, 2022, van der Weerden *et al*., 2013).

Plants use a variety of different mechanisms to resist fungal and oomycete infections (van Esse *et al*., 2020). Anti-microbial peptides (AMPs) are ubiquitous throughout the plant kingdom and play important roles in plant defense against fungal and oomycete pathogens. Their potent anti-fungal and-oomycete activity are vital components of the plant innate immune system (Goyal & Mattoo, 2014, van der Weerden et al., 2013). Defensins, in particular, are one of the largest families of AMPs expressed in plants. Because of their potent antifungal activity, diverse amino acid sequences, and multi-target MoA, plant defensins have emerged as potential antifungal agents for agriculture (Cools *et al*., 2017, Parisi *et al*., 2019, Leannec-Rialland *et al*., 2022). Plant defensins are 45-54 amino acids long, cysteine-rich, cationic peptides characterized by the presence of a cysteine-stabilized αβ motif. This motif consists of one α-helix and three antiparallel β-strands linked by three disulfide bonds. The fourth disulfide bond connects near the N-and C-termini and essentially cyclizes the peptide (Kovaleva *et al*., 2020). The cationic plant defensins exhibit potent broad-spectrum antifungal activity against fungal pathogens *in vitro* and *in planta* (Kaur *et al*., 2011, Kovaleva et al., 2020, Lacerda *et al*., 2014, Leannec-Rialland et al., 2022). We set out to investigate if truncated plant defensins containing the active carboxy-terminal γ-core motif (GXCX3-9C) exhibited potent fungicidal activity and were effective as spray-on fungicides for management of fungal diseases. We further investigated if the smaller defensin-derived peptides exhibited MoA different from those of the full-length parental peptide.

MtDef4 is an evolutionarily conserved 47-amino acid defensin from *Medicago truncatula* that exhibits broad-spectrum antifungal activity against fungal pathogens at low micromolar concentrations. MtDef4 MoA against the plant pathogen *Fusarium graminearum* involves plasma membrane permeabilization, binding to bioactive phosphatidic acid (PA), cell entry, and presumed interaction with intracellular targets (El-Mounadi *et al*., 2016, Sagaram *et al*., 2013, Sagaram *et al*., 2011). Like many other plant defensins, this peptide contains a well-characterized γ-core motif required for its antifungal activity (Sagaram et al., 2013, Sagaram et al., 2011). A chemically synthesized C-terminal 16-amino acid sequence, designated GMA4C, containing this motif and a net charge of +6 inhibited the growth of the fungal pathogen *Fusarium graminearum in vitro* although its potency was four-fold lower than that of MtDef4 (Sagaram et al., 2013). This peptide also exhibited antifungal activity against crown rot fungal pathogens of alfalfa and, in addition, inhibited the growth of plant and human bacterial pathogens *in vitro* (Sathoff *et al*., 2019).

Herein, we explored the potential of the synthetic, MtDef4-derived, structurally less complex, 17-amino acid GMA4CG variants for development as fungicides. We characterized the antifungal activity of several GMA4CG variants generated via amino acid substitutions, C-terminal amidation, and disulfide cyclization that exhibited potent antifungal activity *in vitro* against an aggressive necrotrophic fungal pathogen *Botrytis cinerea*, the causal agent of a gray mold disease in vegetables, fruits, and flowers. In sharp contrast to MtDef4, an interesting characteristic of the GMA4CG variants was that they retained potent antifungal activity in media with elevated salt concentrations. Based on its superior *in vitro* and semi-*in planta* antifungal activity against *B. cinerea*, we focused on GMA4CG_V6 as the lead peptide for *in planta* antifungal activity and MoA studies. GMA4CG_V6 exhibited multi-faceted MoA and rapidly killed fungal cells. When sprayed on *Nicotiana benthamiana* and tomato plants, GMA4CG_V6 conferred strong preventative and curative antifungal activity against *B. cinerea* under the tested conditions.

## RESULTS

### Design of short MtDef4-derived synthetic peptide variants

We based our work on the premise that a small MtDef4-derived peptide spanning the active γ-core motif could serve as a promising template for the design of peptide-based fungicides. We further hypothesized that a small antifungal peptide would be more cell permeant and that with targeted amino acid substitutions of the γ-core motif, we could further enhance its antifungal potency. To test this hypothesis, we generated a truncated set of linear and cyclic variants of the 17-amino acid peptide spanning this motif (Gly30-Cys47s designated GMA4CG_WT and tested them for antifungal activity against *B. cinerea* (Figure 1A).

**Fig 1.**
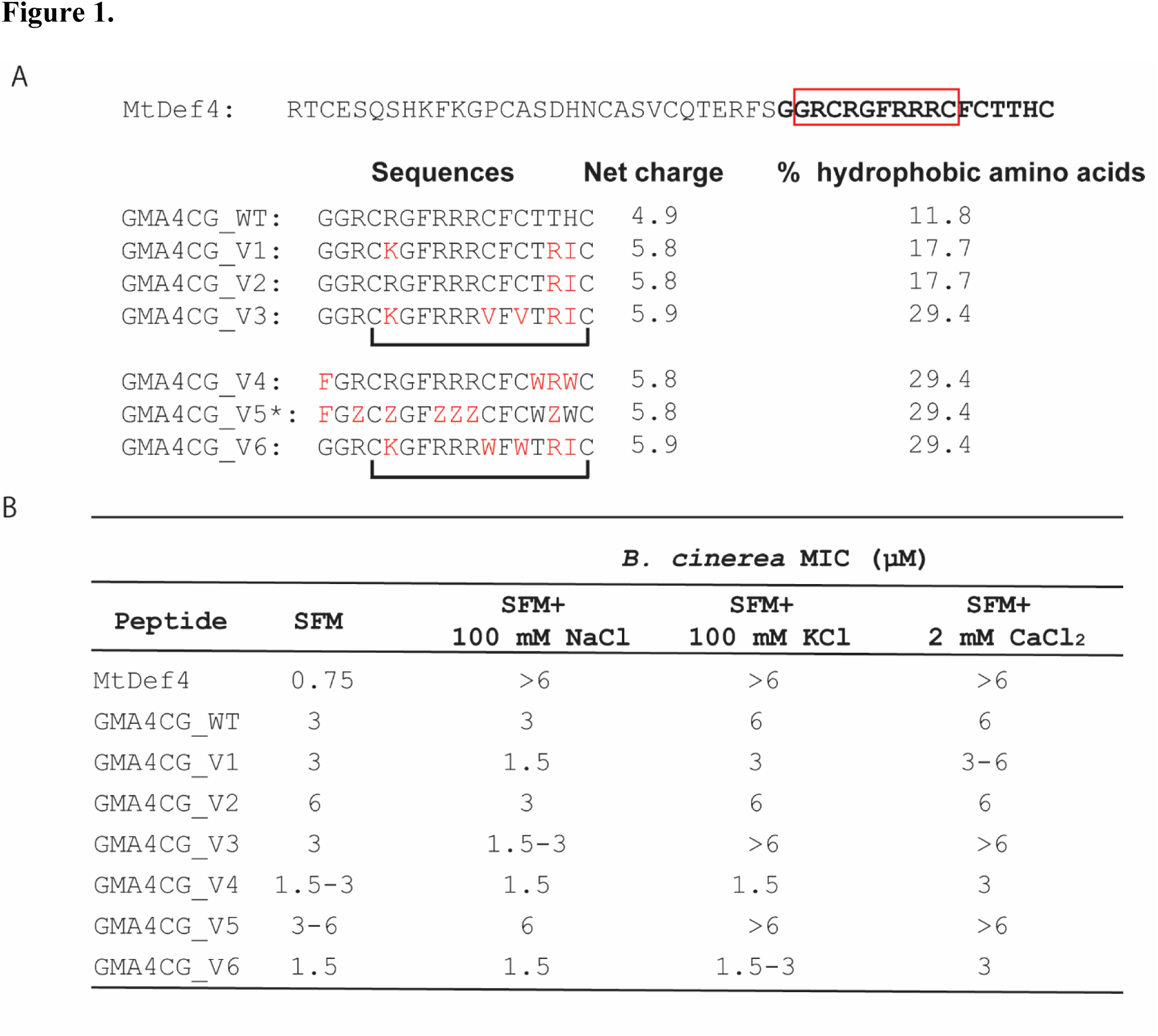
Amino acid sequences and antifungal activity of MtDef4-derived GMA4CG_WT and its variants. **A.** Amino acid sequence, net charge, and percentage of hydrophobic residues for GMA4CG_WT and its variants. The amino acid sequence of MtDef4 is shown on top with its γ-core motif highlighted by a red rectangle. The amino acid substitutions in GMA4CG_WT are shown in red. Disulfide bonds verified in GMA4CG_V3 and GMA4CG_V6 are shown with a link. The letter Z indicates the unnatural amino acid 2, 4-diaminobutyric acid. **B.** The antifungal activity of MtDef4, GMA4CG_WT, and its variants against *B. cinerea* in the absence and presence of cations and against *F. graminearum* in the absence of cations. MIC (Minimal Inhibitory Concentration) of each peptide was determined by the resazurin assay. SFM (Synthetic Fungal Medium) is a low salt medium used in this study.

The substitution of each amino acid with all possible alternative amino acids in GMA4CG_WT was not practical. Thus, our strategy was to substitute cationic or hydrophobic residues in specific positions with the expectation that they would have significant influence on the antifungal activity of this peptide. MtDef4 is an evolutionarily conserved defensin in the plant kingdom and sequences of several homologs from various plants have been reported (Sagaram et al., 2011). GMA4CG_V1 is a natural variant derived from the MtDef4 homolog (KHN09675.1) of *Glycine soja*. This natural variant has three amino acid differences (R5K, T15R and H16I) relative to the sequence of GMA4CG_WT (Figure 1A). GMA4CG_V2 was generated to determine if the substitution of one cationic amino acid with a different cationic amino acid had a notable impact on its antifungal activity. Thus, this variant carried a Lys5 to Arg substitution. We also designed peptide variants (GMA4CG_V3 to GMA4CG_V6) to test the influence of increased hydrophobicity on their antifungal activity (Figure 1A). Thus, each of these variants had 29.4% hydrophobic amino acids as compared with 11.8% in GMA4CG_WT. GMA4CG_WT has four cysteine residues (Cys4, Cys11, Cys13 and Cys17) in its sequence, suggesting a potential to generate a pseudo-cyclic structure via disulfide bond formation between Cys4 and Cys17. Cyclization has been reported previously to increase the antimicrobial activity and selectivity of AMPs (Dathe *et al*., 2004, Tucker *et al*., 2018). Thus, we had cyclic GMA4CG_V3 and GMA4CG_V6 synthesized with a Cys4-Cys17 disulfide bond and replaced Cys11 and Cys13 with either an aliphatic Val (GMA4CG_V3) or an aromatic Trp (GMA4CG_V6) residue. Trp is more effective in interacting with the interfacial regions of lipid bilayers and it promotes antimicrobial activity of peptides (Deslouches *et al*., 2005, Chan *et al*., 2006, Killian & von Heijne, 2000). All peptides were amidated at the C-terminus, a process known to increase the stability of peptides against proteolytic degradation, and therefore, also its biological activity (Lu *et al*., 2020).

### GMA4CG_V6 variants have potent antifungal activity

We assessed the antifungal activity of GMA4CG_WT and its six variants in Synthetic Fungal Medium (SFM) against *B. cinerea in vitro.* GMA4CG_WT inhibited the growth of *B.cinerea* with an MIC value of 3 µM, four-fold greater than that of MtDef4 (Figure 1B). Among all variants, only GMA4CG_V6 displayed two-fold greater antifungal activity than GMA4CG_WT with an MIC value of 1.5 µM (Figure 1B).

To determine if GMA4CG_V6 had fungicidal and/or fungistatic activity, *B. cinerea* conidia were treated with 1.5 µM peptide for 1, 6, and 24 h. Following each treatment, the peptide was removed, and spore germination examined under the microscope. Approximately 85% of the conidia failed to germinate after 1 h of peptide challenge, nearly 100% of the conidia failed to germinate after 6 h, and 100% of the conidia failed to germinate after 24 h of treatment (Figure S1A and B). To assess further the toxicity of GMA4CG_V6, we stained peptide-treated conidial cells with propidium iodide, a cell death marker, and observed that nearly 80% of the conidia stained red within 2 h of peptide treatment indicating fungicidal activity for this peptide (Figure S1C and D).

### The GMA4CG variants, but not MtDef4, retain antifungal activity in the presence of salts

Many naturally occurring AMPs, including plant defensins, are known to lose their antifungal activity in media containing elevated salt concentrations (Kerenga *et al*., 2019, Li *et al*., 2019). We determined the MIC values of GMA4CG_WT and its variants with the R/KGFRRR motif in SFM containing 100 mM NaCl, 100 mM KCl, or 2 mM CaCl2 and compared them to MtDef4. At the greatest concentration tested (6 µM), MtDef4 had no activity against *B. cinerea*. However, GMA4CG_WT and its variants retained their antifungal activity in salt supplemented SFM to varying degrees with GMA4CG_V4 and GMA4CG_V6 retaining the most potent antifungal activity (Figures 1B and S2A). GMA4CG_V6 at 3 µM completely inhibited spore germination in the presence of NaCl and CaCl2 salts (S2B Fig). GMA4CG_V6 was notably more effective than GMA4CG_V3 in the presence of 100 mM KCl and 2 mM CaCl2 (Figures 1B and S2A) indicating that the Val to Trp substitution is important for salt-tolerant antifungal activity in these variants. Collectively, these data revealed that the MtDef4-derived GMA4CG_WT peptide and its variants with a net charge of +5 to +6 exhibited salt-tolerant antifungal activity. In contrast, the parental MtDef4 exhibited salt-sensitive antifungal activity at the same concentrations.

### GMA4CG_WT and GMA4CG_V6 are predominantly random coil structures

The three-dimensional structure of full length MtDef4 contains a γ-core motif composed of β-strands β2 and β3 connected by a positively charged RGFRRR loop (Sagaram et al., 2013). Near-UV CD spectra for GMA4CG_WT and GMA4CG_V6 are shown in Figure S3. Neither spectrum shows features characteristic of a peptide with substantial α-helical (double minima at approximately 222 and 208 nm and an extrapolated maximum around 195 nm) or β-strand (single minimum at around 220 nm and an extrapolated maximum around 195 nm) secondary structure (Greenfield, 2006, Holzwarth & Doty, 1965). The notable difference between the CD spectra of the two peptides was an additional minor minimum at ∼230 nm for GMA4CG_WT and a minor maximum at ∼ 230 nm for GMA4CG_V6. In combination with the major sub-220 nm minimum, the former feature is characteristic of a random coil/unordered structure and the latter feature characteristic of a polyproline II structure suggesting the multiple amino acid substitutions may induce a transition of some random coil to polyproline II structure in GMA4CG_V6 (Lopes *et al*., 2014). Our attempts to predict the structures for GMA4CG and GMA4CG_V6 using the artificial intelligence-based protein structure prediction software AlphaFold (Jumper *et al*., 2021) yielded structures with low confidence. Similarly, *ab initio* structure prediction using the Rosetta protein structure modeling package (Baek *et al*., 2021) did not predict any consistent fold or high-confidence secondary structure elements (data not shown). Hence, the *in silico* modeling data corroborate that experimental CD data, suggesting the peptides have little long-range order and are likely dynamic and disordered in solution.

### The RRRW motif is a major determinant of the salt-tolerant antifungal activity of GMA4CG_V6 with no disulfide bond

A series of linear alanine-substituted variants of our lead peptide GMA4CG_V6 (Figure S4A) were tested to identify amino acids important for antifungal activity. The GMA4CG_V6_Ala3 variant carrying the substitution of RRRW with AAAA lost antifungal activity eight-fold, whereas all other variants lost antifungal activity four-fold (Figures S4A and S4B). In SFM supplemented with 100 mM NaCl, no further loss in the antifungal activity of the variants occurred except in GMA4CG_V6_Ala3, which lost antifungal activity another two-fold (Figure S4C). These results indicated that the RRRW motif is a major determinant of the antifungal activity of GMA4CG_V6 in media supplemented with salts.

### The GMA4CG_V4 and GMA4CG_V6 variants potently reduce gray mold disease symptoms semi-*in planta*

We compared the semi*-in planta* antifungal activity of GMA4CG_WT and its variants using detached leaves of *N. benthamiana* as described (Li et al., 2019). GMA4CG_WT and variants GMA4CG_V1 through GMA4CG_V5 were applied as drops of different concentrations on each leaf with freshly prepared conidial inoculum immediately applied to each drop. Leaves were assessed for attenuation of gray mold symptoms by each peptide 48 h post-inoculation (hpi) relative to no peptide controls by measuring lesion sizes and photosynthetic efficiency (as measured by Fv (variable fluorescence)/Fm (maximal fluorescence)). Increasing concentration of each peptide significantly (*p*≤ 0.0001) reduced lesion sizes (Figures 2A and B). For all peptides, no lesions were formed at 6 µM. At 3 µM, only GMA4CG_V3 and GMA4CG_V4 were effective in suppressing lesion formation. At 1.5 µM, only GMA4CG_V4 completely suppressed disease symptoms. The photosynthetic efficiency data further supported GMA4CG_V4 as being more potent than the other five in reducing gray mold lesions (Figure 2A). We then compared the semi-*in planta* antifungal activity of GMA4CG_V4 and GMA4CG_V6. Both peptides completely suppressed disease symptoms at 1.5 and 3 µM under the applied conditions. However, at a lower concentration of 0.75 µM, GMA4CG_V6 was more effective in suppressing gray mold disease symptoms than GMA4CG_V4 (Figures 2C and D).

**Fig 2.**
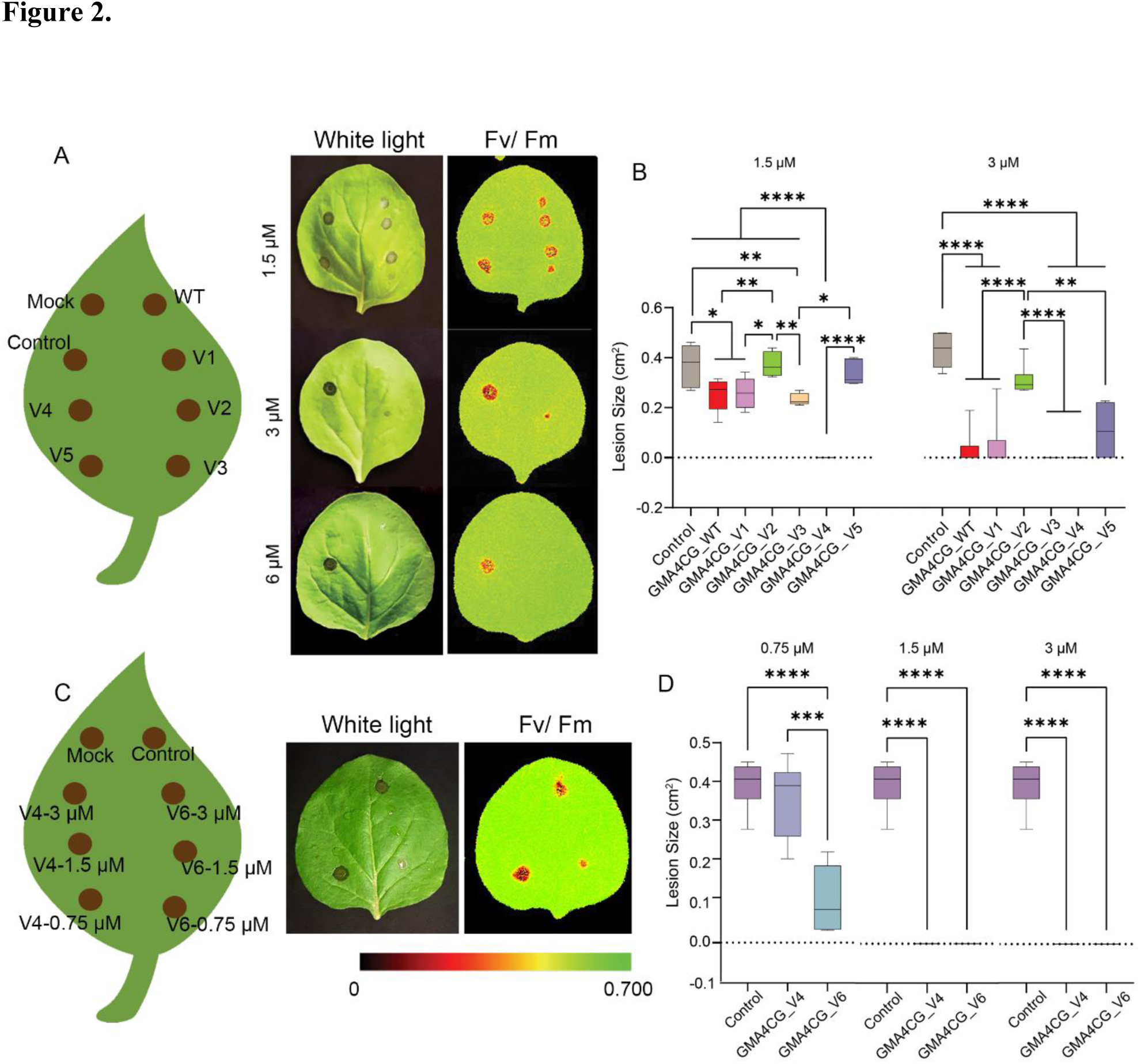
Topical application of GMA4CG_WT and its variants on detached *N. benthamiana* leaves confers resistance to gray mold. **A**. Scheme for application of GMA4CG_WT and GMA4CG_V1-V5 on tobacco leaves. Gray mold lesions on *N. benthamiana* leaves were observed and photographed at 48 hpi. The white light and CropRepoter images of the leaves are shown. **B.** Box & whisker plots were used to represent the quantification of lesion sizes. **C**. Scheme for application of GMA4CG_V4 and GMA4CG_V6 on *N. benthamiana* leaves. The white light and CropReporter images of the infected leaves are shown. **D**. Box & whisker plots were used to represent the quantification of lesion sizes. In **B** & **D**, the horizontal lines represent the median and boxes indicate the 25^th^-75^th^ percentile. One-way analysis of variance (ANOVA) was used to determine statistical differences in lesion sizes between control (*B. cinerea*) and peptides (GraphPad Prism-version 9.4.1). Three technical replications of tobacco leaves (n = 3) were tested in three independent biological replications. Asterisks represent significant differences between different groups (^ns^*p*> 0.05, \**p*≤ 0.05 \*\**p*< 0.01, \*\*\*\**p*≤ 0.0001) using Tukey’s honestly significant difference test for comparison of multiple groups.

### GMA4CG_V6 enhances permeability of the plasma membrane of *B. cinerea* spores and germlings

To determine if GMA4CG_V6 permeabilizes the fungal plasma membrane, the spores and germlings of *B. cinerea* were treated with 1.5 µM GMA4CG_V6 for 30 min followed by the addition of SYTOX^TM^ Green (SG) Nucleic Acid Stain. SG is a membrane-impermeant dye, which fluoresces when it binds to RNA and DNA and is a useful indicator of membrane permeability. SG uptake was observed in nearly all fungal spores and germlings challenged with this peptide, but not in controls without peptide (Figure 3A). Thus, we concluded that one MoA of GMA4CG_V6 involved increased plasma membrane permeabilization.

**Fig 3.**
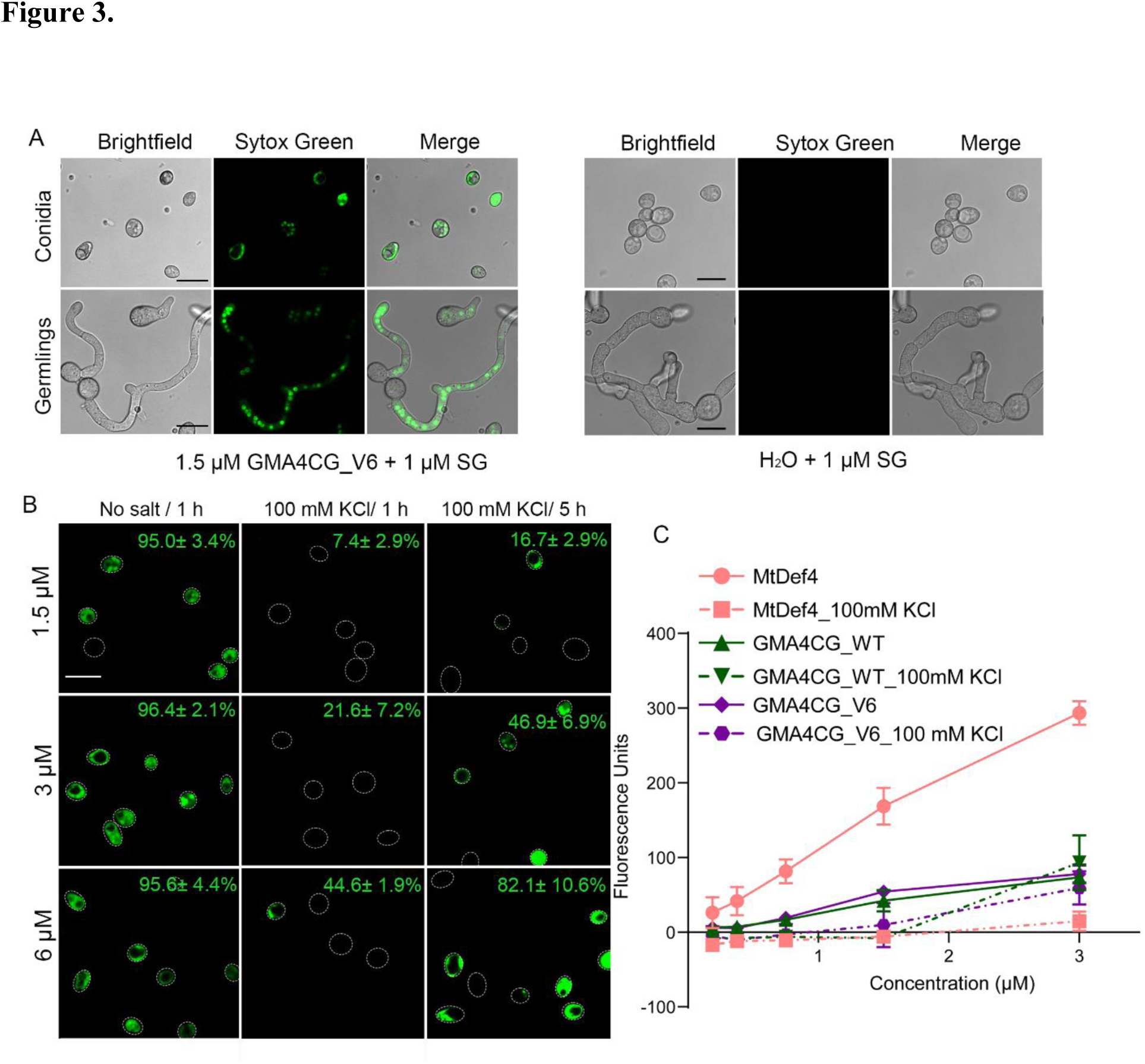
GMA4CG_V6 permeabilizes cell membrane of *B. cinerea*. **A.** Confocal microscopy images of the SG uptake in *B. cinerea* spores and germlings treated with 1.5 µM of GMA4CG_V6 for 30 min. Spores and germlings treated with H2O served as controls. Scale bars = 20 µm. **B.** Representative images showing SG signals in *B. cinerea* spores treated with 1.5, 3, and 6 µM of GMA4CG_V6 for 1 h with no salt, and for 1 h and 5 h with 100 mM KCl. The outline of each spore is highlighted by a white dashed line. Scale bar = 20 µm. The permeabilization rate (%) of each treatment is labeled on the images. **C.** Quantitative measurements of cell membrane permeability in terms of fluorescence emitted by *B. cinerea* germlings treated with various concentrations of MtDef4 (red), GMA4CG_WT (green), and GMA4CG_V6 (purple) with (dashed line) and without (solid line) 100 mM KCl. Membrane permeabilization increased with increasing concentration of each peptide. Values are means of two biological replications. Error bars indicate standard deviations. Fluorescence measurements were taken at 5 h.

Since antifungal activity of GMA4CG_V6 was cation-tolerant, we determined if membrane permeability still occurred in the presence of 100 mM KCl. In the absence of this salt, 95% of the conidia exposed to 1.5 µM GMA4CG_V6 for 1 h took up SG, indicating increased membrane permeability. No further time-or concentration dependent increase in membrane disruption was observed in the absence of this salt (Figure 3B). By contrast, uptake of SG occurred in less than 10% of the conidia treated with 1.5 µM GMA4CG_V6 for 1 h in the presence of 100 mM KCl. Even after 5 h, only ∼17% of the conidia took up SG. A high concentration of 6 µM GMA4CG_V6 was needed for the permeabilization of ∼80% of conidia after 5 h (Figure 3B). Thus, the membrane-permeabilization ability of GMA4CG_V6 was negatively impacted by the presence of cations; however, the peptide was still effective in permeabilizing the fungal plasma membrane at increased concentrations with longer exposure.

We also used our SG assay to compare the ability of GMA4CG_V6, GMA4CG_WT, and MtDef4 to permeabilize the plasma membrane in germlings in the absence and presence of 100 mM KCl. In the absence of KCl, MtDef4 was more effective in permeabilizing the plasma membrane than GMA4CG_WT or GMA4CG_V6. However, in the presence of this salt, MtDef4’s ability to permeabilize the plasma membrane was completely lost at low concentrations (< 3 µM). In contrast, GMA4CG_WT and GMA4CG_V6 retained their ability to permeabilize the plasma membrane at < 3 µM (Figure 3C).

### GMA4CG_V6 is internalized into the vacuole and cytoplasm of B. cinerea in a time and concentration dependent manner

Tetramethyl rhodamine (TMR)-labeled GMA4CG_V6 was imaged using time-lapse confocal microscopy to gain insight into the temporal nature of peptide internalization in *B. cinerea* germlings. At an MIC of 1.5 µM, the labeled peptide first bound to the cell surface within the first 3 min and started translocating into vacuoles from multiple entry foci by 19 min (Figure 4A). At this time point, bright field images of germlings showed abnormal morphology with vacuoles becoming visible. At 27 min, the vacuoles began to expand by coalescence of small vacuoles (Figure 4A, arrows) and the peptide localized into the cytoplasm. Shortly after the peptide’s appearance in the cytoplasm the vacuoles appeared to lyse with the peptide rapidly dispersing throughout the germling cytoplasm and inducing cell death (Fig 4A, T= 59).

**Fig 4.**
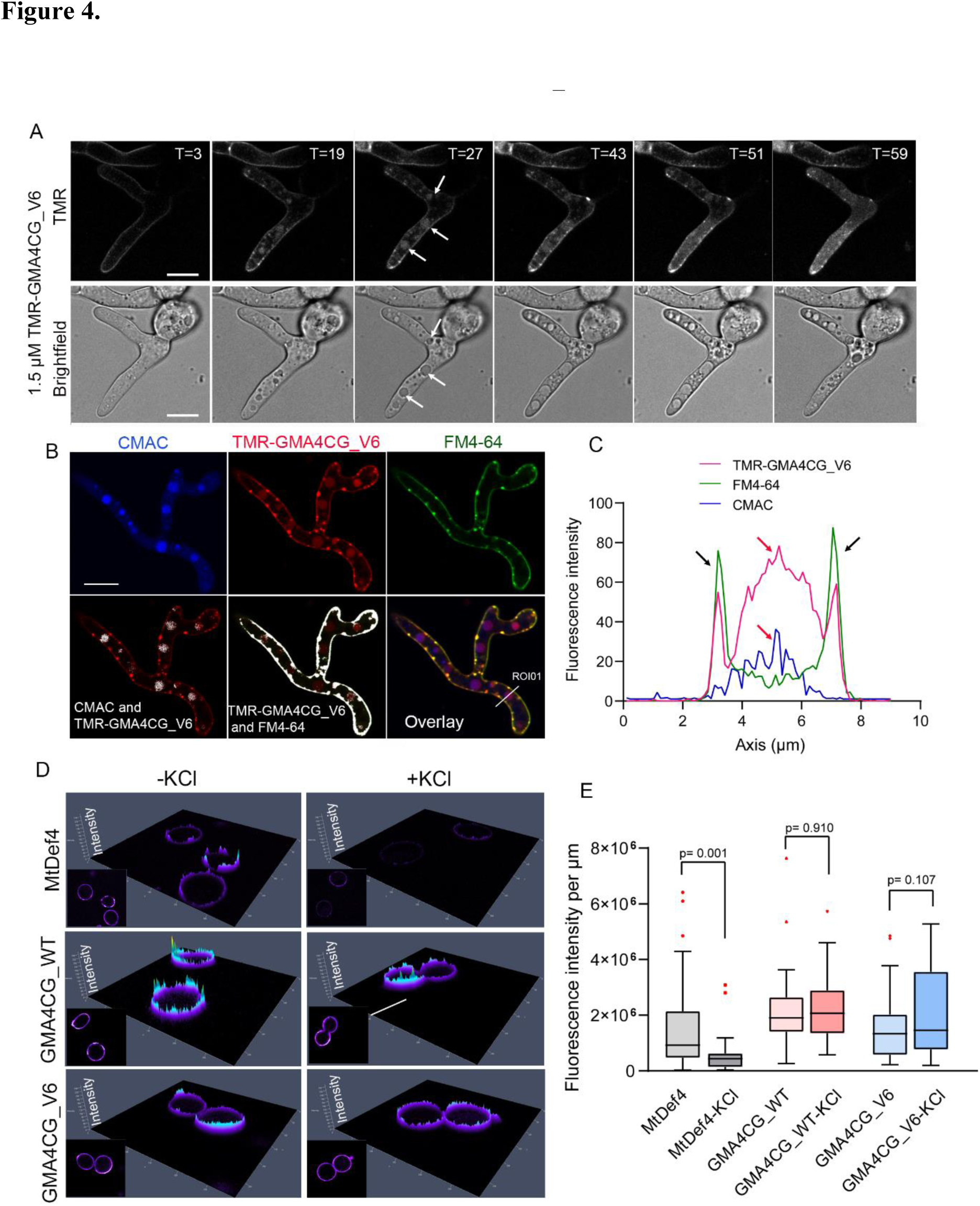
Internalization of GMA4CG_V6 into *B. cinerea*. **A.** Time-lapse confocal microscopy images of *B. cinerea* showing internalization of 1.5 µM TMR-GMA4CG_V6 over time (T in min). Images were captured every 4 min starting from 3 min. Arrows indicate vacuoles. Scale bars = 10 µm. **B.** Individual fluorescent (CMAC, TMR, and FM4-64) and channel merged images of *B. cinerea* germlings treated with GMA4CG_V6 for 30 min. The overlap signal of CMAC and TMR or TMR and FM4-64 are shown as white pixels. The threshold for each channel and the background was 50% and 20%, respectively. Scale bars = 10 µm. **C.** The fluorescent signal intensity of the ROI in **B**. Black arrows indicates TMR-GMA4CG_V6 and FM4-64 signals overlap at the membrane and the red arrows indicate TMR-GMA4CG_V6 and CMAC signals overlap at the vacuoles. **D.** Cell surface fluorescence shown in 2.5D. The vertical direction indicates the fluorescence intensity of each pixel with taller peaks representing greater fluorescence intensity. The corresponding 2D images are shown. **E.** Box plot for cell surface fluorescence quantification. Total fluorescence intensity of each cell was normalized to the cell perimeter length. The lower threshold was set to 7000, the substrate signal in the cytoplasm. Statistical analyses were performed using a two-tailed *t*-test assuming unequal variance (GraphPad Prism 8.0). *P*-value over 0.05 indicated no significant difference and less than 0.05 indicates a significant difference. n = 24-39.

At a sub-lethal concentration of 0.5 µM, the labeled peptide was localized at the germling periphery and formed presumptive endosomes in the cytoplasm. No further progression of the peptide in the germling cells occurred during the 1 h challenge (Figure S5A). At high concentration (20 µM), TMR-labeled GMA4CG_V6 was internalized by germlings as early as 4 min. The peptide rapidly accumulated into the germling cytoplasm and within 13 min was completely dispersed throughout the cell’s interior (Figure S5B). This suggests that at high concentrations, GMA4CG_V6 may largely bypass intermediate endosome or vacuole localization and accumulate directly in the cytoplasm, or highly accelerate the translocation from vacuole to cytoplasm and finally accelerate cell death.

The subcellular localization of TMR-GMA4CG_V6 was determined using two dyes which specifically stain the plasma and endosomal membranes (FM4-64) and vacuole lumen (CMAC). Within 30 min of challenging germlings with peptide, TMR-GMA4CG_V6 co-localized with CMAC in vacuoles and with FM4-64 at the plasma membrane and entry foci (Figure 4B, highlighted white). Based on the region of interest (ROI01) in Figure 4B, the fluorescence intensity curve clearly showed that the peak signal of TMR-GMA4CG_V6 overlapped with the signal of FM4-64 in membrane and CMAC in vacuole (Figure 4C). To determine whether GMA4CG_V6 bound to the cell wall of *B. cinerea*, a cell wall specific dye Calcofluor White (CFW) was incubated with TMR-GMA4CG_V6 treated *B. cinerea* germlings. As illustrated in Figures S6A and B, CFW overlapped with TMR fluorescent signal in images captured by confocal microscopy. Considering the thickness and proximity of germling cell wall and plasma membrane and the limitation of the resolution of the confocal microscopy, super-resolution microscopy (specifically, lattice structured illumination microscopy (SIM) was used to further evaluate co-localization of TMR-GMA4CG_V6 with the cell wall. The fluorescent signal of CFW relative to TMR showed the TMR signal predominantly just inside the cell wall (Figures S6C, D and E), as shown by arrows, indicating that dominant TMR-GMA4CG_V6 signal was consistent with intense plasma membrane localization at the cell periphery, although low level cell wall binding could not be ruled out.

To further study the binding affinity of GMA4CG_V6 to specific subcellular components in *B. cinerea* germlings, a series of Fluorescence Recovery After Photobleaching (FRAP) experiments were performed (Figure S7A–C). We specifically studied the mobility of TMR-GMA4CG_V6 to subcellular components including germling cell periphery, vacuoles, and cytoplasm. In our previous study, a significant difference in the mobility of the antifungal peptide NCR044 to germling and conidial cell walls was observed (Velivelli *et al*., 2020a), thus, we performed FRAP experiments with conidial spore periphery as well. According to the normalized fluorescence recovery curve, TMR-GMA4CG_V6 recovery in the germling cytoplasm had the greatest overall mobility among all four targeted components and the mobility in vacuoles was relatively low (Figure S7A). Immobile fraction and half time of fluorescence recovery data were also collected as summarized in Figures S7B and C. GMA4CG_V6 in germling vacuoles showed a significantly higher (*p*< 0.0001) immobile fraction (82.6 ± 3.2%) than other subcellular components and the recovery time was much shorter (Tau= 19.27 sec, R^2^= 0.8496). In cell periphery of germlings and conidia, no significant difference was observed in the immobile fraction (52.5± 4.2% and 51.0 ± 4.9%, respectively, *p*> 0.05) or in the half recovery time (37.4 sec, R^2^= 0.9133 and 38.4 sec, R^2^= 0.9257, respectively). The immobile fraction of the peptide was lower (*p*= 0.0031) in germling cytoplasm (35.2± 4.3%) as compared to that in germling cell periphery and vacuole but showed no significant difference with that of the spore periphery. The mobility of peptide in germling cytoplasm was the highest among the four selected subcellular components (Tau= 40.68 and R^2^= 0.9636). Collectively, the FRAP analysis indicated that the mobility of the peptide at the cell periphery and cytoplasm was similar but much different in the vacuoles. The low mobility in vacuoles is likely because transport to the vacuole lumen is not a continuum like other compartments and multistep mobility into the vacuole itself would be very high as the Tau values show.

Because of the difference in the cation tolerant antifungal activity of MtDef4 and GMA4CG, we examined the localization of MtDef4, GMA4CG_WT, and GMA4CG_V6 on the spores of *B. cinerea* in the presence of 100 mM KCl. The fluorescently labeled MtDef4 localized to the cell periphery in the absence of this salt, but this localization was significantly reduced in its presence (*p* = 0.001). In contrast, cell periphery localization of GMA4CG_WT and GMA4CG_V6 was not affected by salt (*p*> 0.05) (Figures 4D and E).

### GMA4CG_WT and GMA4CG_V6 bind to multiple phosphoinositides

MtDef4 binds to phosphatidic acid (PA) in fungal membranes as part of its MoA (18). Therefore, we tested if MtDef4-derived GMA4CG_V6 also binds to PA. A protein-lipid overlay assay was performed with various biologically relevant phospholipids immobilized on a lipid strip (Echelon Bioscience, UT). GMA4CG_V6 bound to several phosphoinositides including phosphatidylinstol (PI) mono-and bisphosphates, PA, and phosphatidylserine, but with higher affinity for PI4P and PI5P than for PA (Figure S8). GMA4CG_WT also bound to the same phospholipids as GMA4CG_V6. His6, the C-terminal tag attached to each peptide used in the assay for antibody detection, did not bind to the tested phospholipids. These results indicated that GMA4CG_V6 was more promiscuous in its binding to the membrane resident phospholipids than MtDef4 (Sagaram et al., 2013).

### Spray-applied GMA4CG_V6 provided preventative and curative antifungal activity against the gray mold disease

We tested the potential of GMA4CG_WT and GMA4CG_V6 to provide preventative protection from gray mold disease by spraying two-week-old *N. benthamiana* plants with each peptide at concentrations of 12 and 24 µM followed 2 h later with the spray-inoculum of *B. cinerea* spores. Each peptide spray-applied at 12 or 24 µM reduced gray mold disease symptoms compared with control plants inoculated with pathogen only (Figures 5A and S9A). Both peptides significantly prevented the spread of gray mold on *N. benthamiana* plants (Figures 5A and 5B). Notably, plants sprayed with GMA4CG_V6 and subsequently inoculated with the pathogen looked as symptom free as non-inoculated mock (water)-sprayed plants. This data clearly demonstrated higher potency of GMA4CG_V6 over GMA4CG_WT in reducing gray mold disease in *N. benthamiana* plants in a preventative manner. In addition, GMA4CG_V6 was more effective in limiting disease severity than GMA4CG (Figures S9A and B).

**Fig 5.**
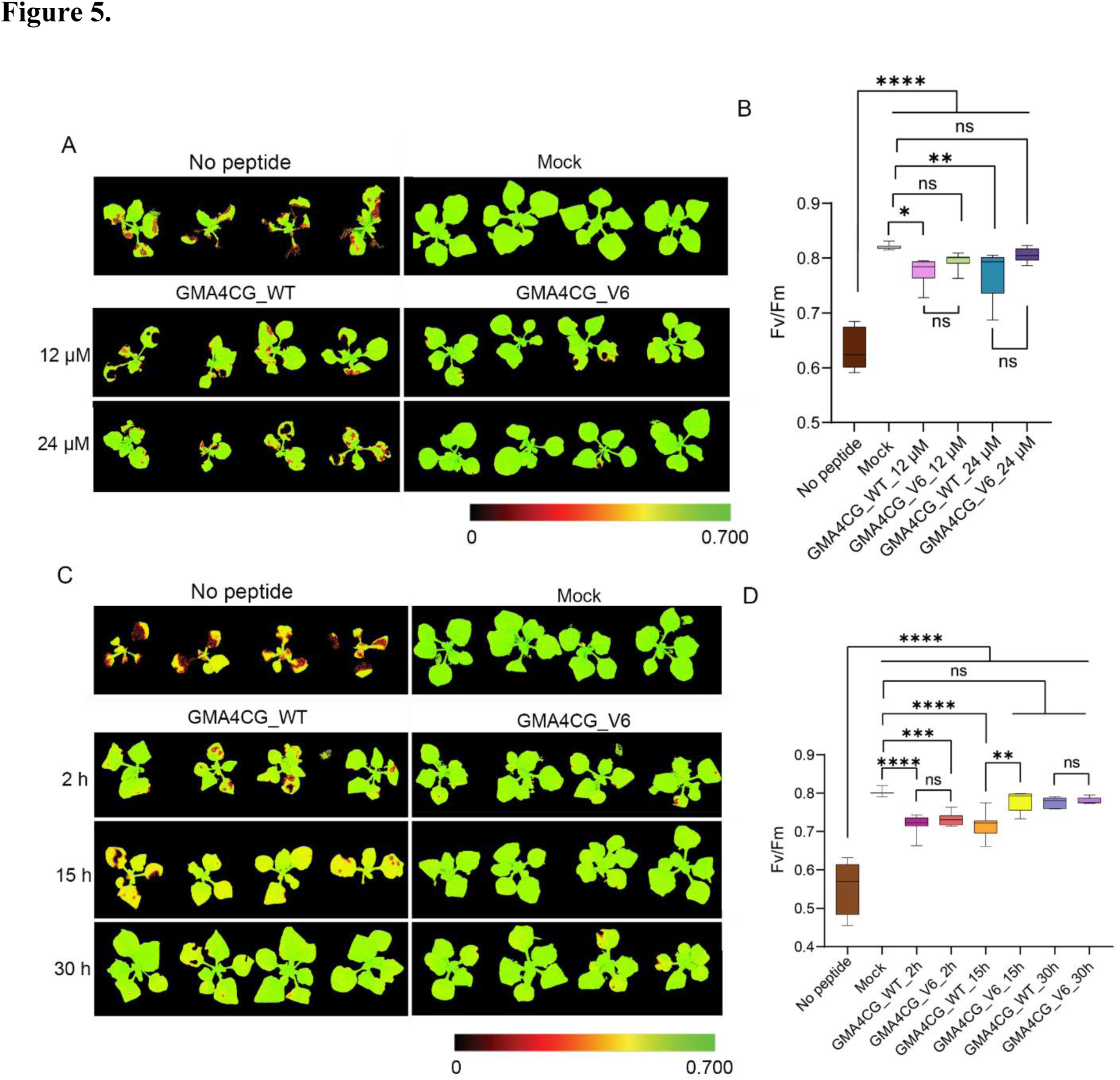
GMA4CG_V6 exhibits preventative and curative antifungal activity against gray mold in *N. benthamiana* plants. **A.** Preventative activity of GMA4CG_V6. Four-week-old *N. benthamiana* plants were sprayed with either 2 mL of water or peptide (GMA4CG_WT or GMA4CG_V6) at 12 µM or 24 µM concentrations prior to spray-inoculation with 1 mL of 8 × 10^4^ spores per mL of *B. cinerea* fresh spore suspension. The mock plants were only sprayed with water (no fungal spore suspension). The CropReporter images of plants were taken 72 hpi. Maximum quantum yield of PSII photochemistry (Fv/Fm) was determined at 72 hpi. Red color represents class I, low photosynthesis efficiency attributed to extensive tissue damage. Green color indicates class V, high photosynthesis efficiency. **B.** The calculated photosynthetic quantum yield (Fv/Fm) for individual *N. benthamiana* plants. **C.** Curative antifungal activity of GMA4CG_V6. Four-week-old *N. benthamiana* plants were first spray-inoculated using 1 mL of 8 × 10^4^ *B. cinerea* spore suspension. At 2, 15, or 30 hpi the plants were sprayed with 2 mL of peptide (GMA4CG_WT or GMA4CG_V6) at a concentration of 12 µM or water. The mock plants were only sprayed with water (no fungal spore suspension). Maximum quantum yield was determined as described above. **D.** The calculated photosynthetic quantum yield (Fv/Fm) for individual *N. benthamiana* plants. In **B** and **D** the horizontal lines represent the median and boxes indicate the 25^th^ and 75^th^ percentiles. One-way analysis of variance (ANOVA) was used to determine statistical difference between control (*B. cinerea* alone), mock (water alone), and peptide-treated inoculated plants (GraphPad Prism-version 9.4.1). Each technical replication included four plants (n = 4). Three biological replications were performed. Asterisks represent significant differences between different groups with *p* values (^ns^*p*> 0.05, \**p*≤ 0.05 \*\**p*< 0.01, \*\*\*\**p*≤ 0.0001) determined using Tukey’s honestly significant difference test for comparison of multiple groups.

We also tested GMA4CG_WT and GMA4CG_V6 for their ability to provide curative protection from this disease at three different time points following fungal infection. *N. benthamiana* plants were pre-inoculated with freshly prepared *B. cinerea* spores for 2, 15, and 30 h after which each peptide at 12 µM was sprayed onto the plants. *B. cinerea* spores germinate after 12 h (Bi *et al*., 2021), and thus, fungal infection of the leaf surface was already initiated at 15 and 30 hpi. Disease symptoms were rated at 72 hpi. Both peptides provided significant curative (*p* = 0.0019) protection from gray mold (Figures 5C and S9C). However, GMA4CG_V6 conferred significantly greater (*p*<0.0001) curative antifungal activity than GMA4CG_WT sprayed on plants pre-inoculated for 15 h (Figure 5D). The reduced disease severity in GMA4CG_V6 sprayed plants further confirmed greater efficacy of this peptide (Figure S9C). In plants pre-inoculated for 2 h or 30 h, both peptides provided robust control of disease symptoms with similar efficacy (Figures 5D and S9D). Furthermore, our data showed the potential of GMA4CG_V6 in reducing the spread of disease symptoms (Figure S9D).

Next, we tested the curative antifungal activity of GMA4CG_WT and GMA4CG_V6 on tomato (cv. Mountain Spring) plants pre-inoculated for 12 h and subsequently sprayed with each peptide at a concentration of 6, 12, and 24 µM. Both peptides conferred curative antifungal activity at all three concentrations (Figure 6A). At concentrations of 12 and 24 µM, no significant difference was observed between the two peptides in their ability to reduce disease symptoms. Both peptides provided robust curative protection with pathogen-challenged plants appearing similar to water-sprayed non-inoculated mock plants. However, at a concentration of 6 µM, GMA4CG_V6 conferred significantly greater (*p*< 0.0001) curative antifungal activity than GMA4CG_WT (Figure 6B).

**Fig 6.**
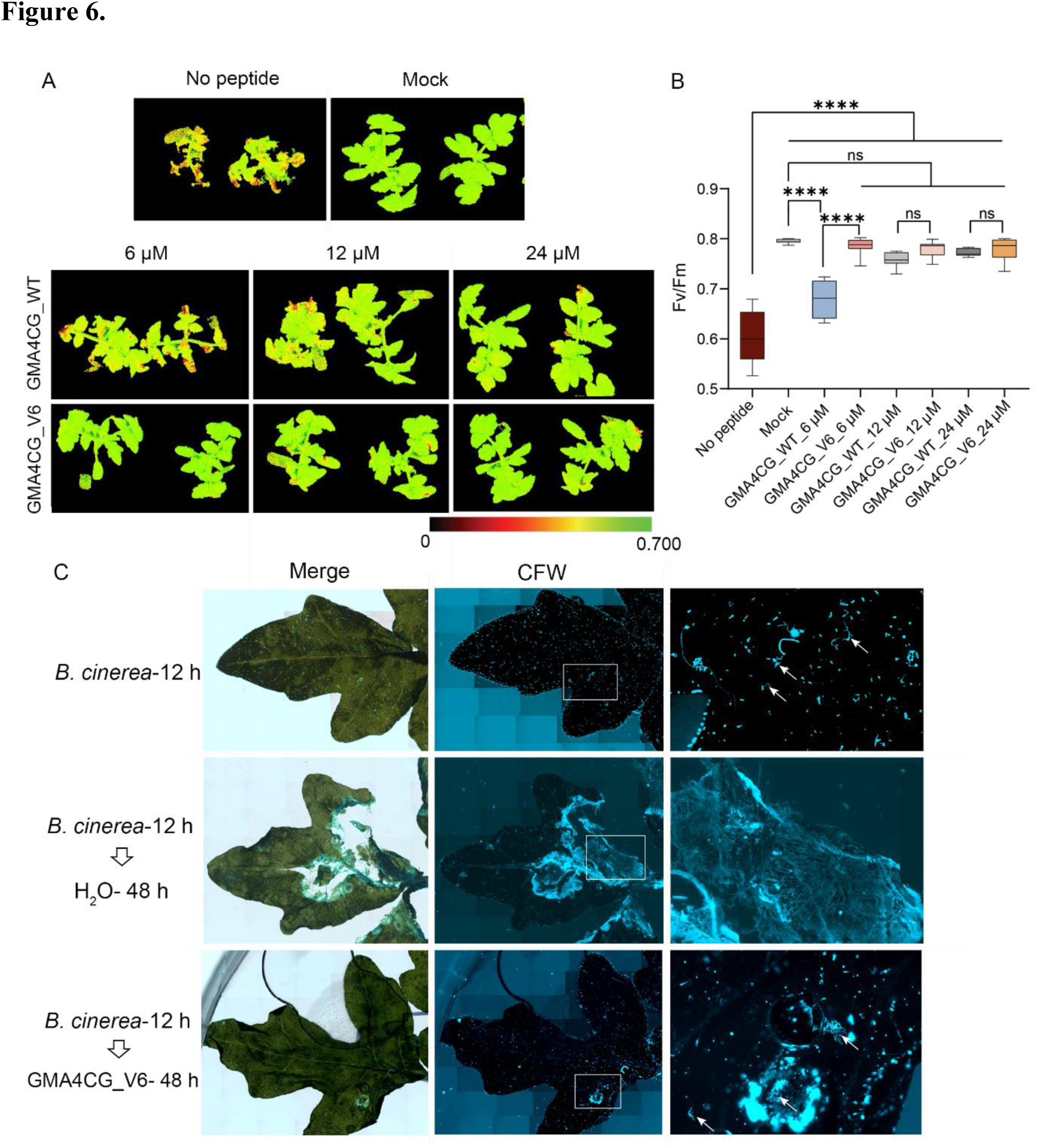
GMA4CG_V6 exhibits curative antifungal activity against gray mold in tomato plants. **A.** Three-week-old tomato plants were spray-inoculated with 1 mL of 8 × 10^4^ *B. cinerea* spore suspension. At 12 hpi, plants were sprayed with 2 ml water or 2 ml peptide (GMA4CG_WT or GMA4CG_V6) at concentrations of 6, 12, or 24 µM. The CropReporter images of the plants were taken 72 hpi and the maximum quantum yield of PSII photochemistry (Fv/Fm) was determined. Red color represents class I, low efficiency photosynthesis attributed to extensive tissue damage. Green color represents class V, high efficiency photosynthesis. **B.** The calculated photosynthetic quantum yield (Fv/Fm) for individual tomato plants. The horizontal lines represent the median and boxes indicate the 25^th^ and 75^th^ percentiles. One-way analysis of variance (ANOVA) was used to determine statistical difference between control (*B. cinerea* alone), mock (water alone), and peptide-treated inoculated plants (GraphPad Prism-version 9.4.1). Each technical replication included four plants (n= 4). Three biological replications were performed. Asterisks represent significant differences between different groups with *p* values (^ns^*p*> 0.05, \**p*≤ 0.05 \*\**p*< 0.01, \*\*\*\**p*≤ 0.0001) determined using Tukey’s honestly significant difference test for comparison of multiple groups. **C.** Germination and growth of *B. cinerea* spores on the tomato leaf surfaces in the presence of GMA4CG_V6 or water. Tomato plants were inoculated with *B. cinerea* spores. Infection was allowed to progress for 12 h and then the plants were sprayed with water or with GMA4CG_V6. Forty-eight hours post peptide or water treatment, leaf samples were fixed in 4% paraformaldehyde and stained with Calcofluor White for 30 min. ZEISS Axio ZoomV16 was used to capture images. Arrowheads point to the germinated hyphae of *B. cinerea*.

Visual rating of disease severity clearly revealed superiority of GMA4CG_V6 in controlling gray mold disease in tomato plants (Figures S10A and B). The cell wall staining of the tomato leaves from control plants infected with the pathogen alone and from infected plants sprayed with GMA4CG_V6 confirmed strong inhibition of fungal growth provided by this peptide (Figure 6C).

### GMA4CG_V6 binds strongly to the tomato leaf surface and is not internalized into plant cells

To determine the distribution of GMA4CG_WT and GMA4CG_V6 on leaf surfaces, fluorescently labeled peptide was applied onto detached tomato leaves and the uptake of each peptide by the plant cells was followed using confocal microscopy. In leaves with no applied peptide, only autofluorescence signal from the plant cells was observed (Figure S11A). Both GMA4CG_WT and GMA4CG_V6 bound to the exterior of plant cell walls, but no internalization into plant cells was detected (Figure S11B). After washing the leaves with water, the fluorescence intensity of GMA4CG_WT was significantly reduced (*p*< 0.0001) but that of GMA4CG_V6 was only slightly reduced (*p*= 0.1146) (Figure S11C). This result suggests that GMA4CG_V6 adheres to the tomato leaf surface more than GMA4CG_WT.

## DISCUSSION

Modern agriculture relies heavily upon systemic single-site fungicides with preventative and curative antifungal properties. However, rapid evolution of fungal resistance to fungicides poses a major threat to crop protection and food security worldwide. New environmentally friendly antifungal compounds with multifaceted MoA are needed to mitigate resistance development. Here, inspired by the broad-spectrum antifungal activity, structure, and multi-faceted MoA of the plant defensin MtDef4, we explored the potential of a short 17-residue peptide for development as a peptide-based fungicide.

We designed and synthesized six 17-amino acid GMA4CG variants spanning the active γ-core motif of a plant defensin MtDef4 (El-Mounadi et al., 2016, Sagaram et al., 2013, Sagaram et al., 2011). We made several modifications of the GMA4CG peptide that maintained or enhanced its antifungal activity against an economically important fungal pathogen *B. cinerea*. Although we could not perform complete analysis of all amino acid substitutions, our selected changes revealed the importance of cationic and hydrophobic residues for antifungal activity of this peptide. Amino acids we chose to replace were guided partly by the corresponding amino acid sequences of the homologs of MtDef4 from other plants. In addition, replacement of the internal Cys11 and Cys13 residues to the aromatic hydrophobic amino acid Trp enhanced antifungal activity. The importance of Trp at these positions was revealed by our observation that an aliphatic hydrophobic residue Val at these positions was less effective. It is likely that interactions of the two Trp residues with the Arg residues present in the GMA4CG_V6 primary amino acid sequence contribute significantly to peptide stability (Shi *et al*., 2002). Further, Trp is known to insert into membranes containing anionic phospholipid head groups with high efficiency, thus contributing to the increased fungal membrane permeability by this designed peptide (Schibli *et al*., 2002).

We synthesized GMA4CG_V6 as a cyclic peptide with a disulfide bond connecting Cys4 and Cys17 with the notion that it would be more stable, more fungal cell permeable, more readily internalized into fungal cells, and confer higher antifungal activity than its linear counterpart. Cyclic peptides are touted as promising antimicrobial molecules since they are more stable to proteolytic degradation, have high affinity and selectivity for targets, and better cell membrane permeability (Falanga *et al*., 2017).

GMA4CG_WT and GMA4CG_V6 each have random coil/unordered structures as determined experimentally by steady-state wavelength CD spectra showing the absence of any significant α-helical or β-strand secondary structure content (Greenfield, 2006, Holzwarth & Doty, 1965). Both peptides are predominately random coil in nature with a suggestion of some polyproline II structure in GMA4CG_V6 (Lopes et al., 2014).

Antifungal activity of several cationic plant defensins is abrogated in well-defined synthetic fungal growth medium containing elevated levels of Na^+^ or Ca^2+^ (Spelbrink *et al*., 2004, Terras *et al*., 1992, Kerenga et al., 2019, Li et al., 2019). At the highest concentration of 6 µM tested, MtDef4 exhibited no antifungal activity against *B. cinerea* in medium containing elevated levels of Na^+^, K^+^, or Ca^2+^. The presence of cations likely weakens the electrostatic interactions between this cationic plant defensin and negatively charged fungal membranes. Surprisingly, all six truncated peptide variants derived from MtDef4 exhibited antifungal activity against *B. cinerea* in fungal growth medium containing elevated levels of Na^+^, K^+^, or Ca^2+^. Among the six variants, GMA4CG_V6 was the most potent antifungal agent. While the molecular basis for cation-tolerant antifungal activity of these peptides is unclear, the RRRW motif clearly plays an important role in conferring cation-tolerant antifungal activity.

The structure and MoA of GMA4CG_V6 are unlike those of any fungicide currently commercially available. The preventative and curative plant protection properties of GMA4CG_V6 is likely due to its ability to kill fungal pathogens rapidly. It does not have a discrete intracellular molecular target, but instead interferes with multiple cellular processes (Figure 7). A major characteristic of antifungal peptides is their ability to permeabilize the plasma membrane of fungal pathogens. GMA4CG_V6 quickly binds to the cell surface of fungal spores and germlings and subsequently disrupts the plasma membrane of these cells near the entry points. The co-localization of GMA4CG_V6 with the vacuolar dye CMAC revealed that this peptide, at MIC values, localizes to the plasma membrane and vacuoles in germlings of *B. cinerea*. However, it remains to be determined if vacuole localization is required for the antifungal activity of this peptide, or simply a fungal detoxification step. It is of interest to note that a small rationally designed antifungal hexapeptide PAF26 with the sequence Ac-RKKWFW-NH2 also localizes to vacuoles in the model fungus *Neurospora crassa* (Munoz *et al*., 2012). Like PAF26, GMA4CG_V6 at MIC enters vacuoles and causes their coalescence and lysis. It subsequently accumulates in the cytoplasm. At elevated concentration, GMA4CG_V6 appeared to accumulate rapidly in the cytoplasm. MtDef4 binds to the bioactive phospholipid PA as part of its MoA (El-Mounadi et al., 2016, Sagaram et al., 2013). In contrast, GMA4CG_V6 derived from this defensin binds to multiple bioactive phospholipids including PA. It preferentially binds to the phosphoinositides PIPs in the phospholipid-protein overlay assay. Further studies are needed to understand the role promiscuous interactions with multiple phospholipids play in the multi-faceted MoA of GMA4CG_V6.

**Fig 7.**
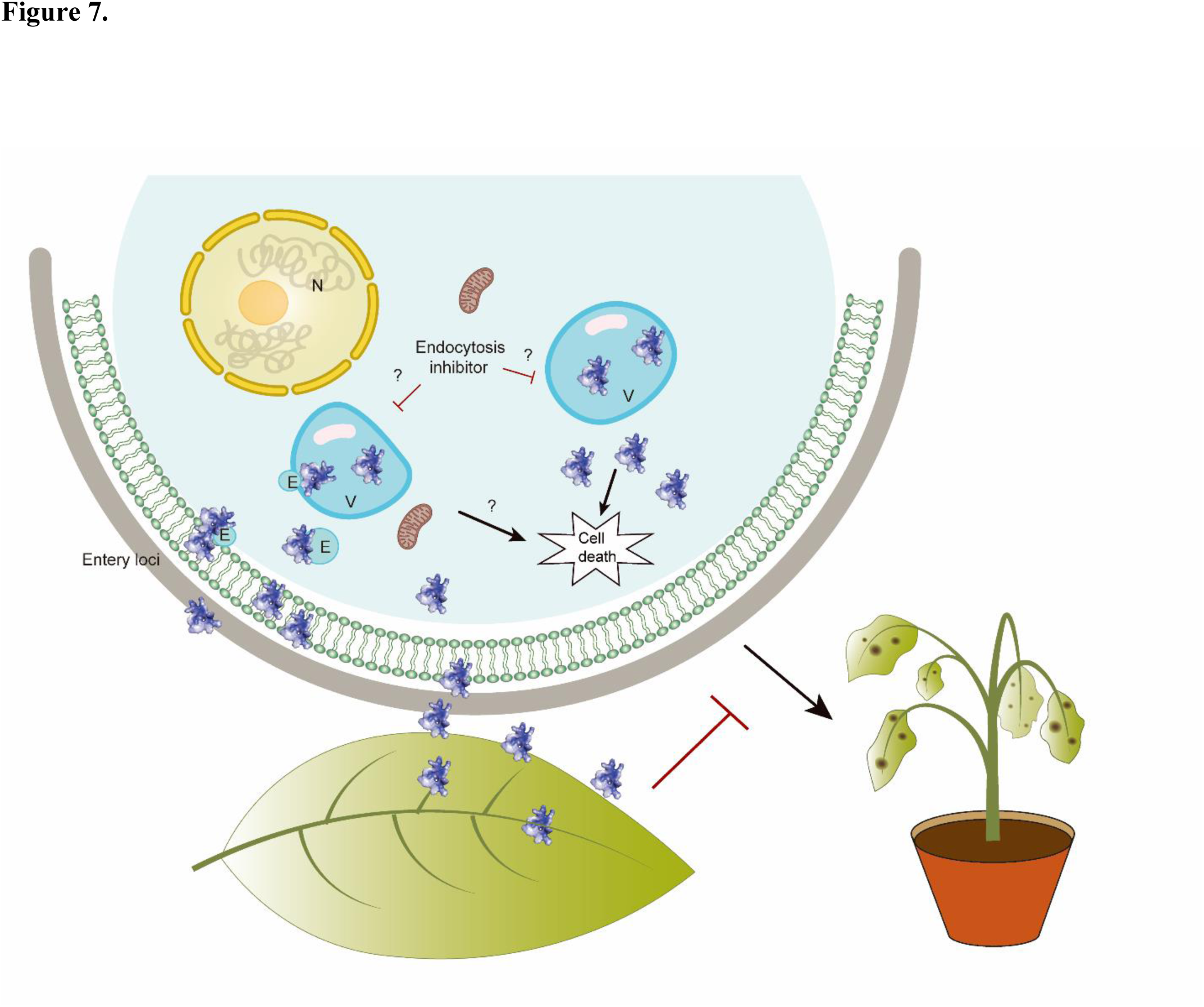
A proposed model for the modes of antifungal action of GMA4CG_V6 against *B. cinerea*. GMA4CG_V6 sprayed on *B. cinerea* infected leaves first binds predominantly to the fungal plasma membrane with little or no binding to the cell wall. Binding to plasma membrane-resident phosphoinositides presumably plays a role in membrane permeabilization. At low concentration, GMA4CG_V6 is internalized into cell vacuoles. It remains to be determined if this peptide is internalized into vacuoles through endocytic mechanisms. At high concentration, GMA4CG_V6 targets the cytoplasm immediately following cell entry. Cell death occurs only after GMA4CG_V6 diffuses into the cytoplasm and not while in vacuoles. N: nucleus, V: vacuole, E: endosome.

Current crop protection research is guided by a strong demand for sustainable fungicides that are efficacious, less prone to resistance development, environmentally friendly, and nonphytotoxic. In semi-*in planta* antifungal assays using detached *N. benthamiana* leaves, all six GMA4CG analogs significantly reduced gray mold disease lesions caused by *B. cinerea* infection with GMA4CG_V6 proving most effective. Under the conditions employed, spray application of GMA4CG_V6 conferred strong preventative and curative antifungal activity against this disease in young *N. benthamiana* and curative antifungal activity in tomato. These fungicidal effects were observed readily even though the dose, application frequency, and distribution formulation have yet to be optimized. In our studies, GMA4CG_V6 was more effective than GMA4CG_WT in reducing disease symptoms suggesting that the amino acid substitutions introduced in GMA4CG_V6 are important for its *in vitro* as well as *in planta* antifungal activity. Although defensin-derived peptides offer the benefits of small size, structural simplicity, and efficiency of peptide modifications, their commercial use as peptide fungicides will require new innovations in peptide production at scale to be economical. In addition, efficacy of these peptides for fungal disease control will need to be assessed in large-scale field trials and their stability to proteolytic degradation and potential phytotoxicity under field conditions will also need to be evaluated.

Recent studies have shown the promise of antifungal peptides as topical fungicides. A fusion of the animal AMP dermaseptin and thanatin inhibits the germination of the spores of the obligate biotrophic fungal pathogen *Phakopsora pachyrhizi* and reduces the symptoms of Asian rust when spray-applied to soybean leaves (Schwinges *et al*., 2019). Spray-application of the antifungal peptide NCR044 from *M. truncatula* on tomato and *N. benthamiana* plants provided protection from the gray mold disease (Velivelli et al., 2020a). The present study showed that a designed defensin-derived peptide spanning the functionally active γ-core motif with multi-faceted MoA exhibited improved preventative and curative antifungal activity against the gray mold disease of plants. Hundreds of sequences divergent antifungal plant defensins in the AMP database can now be exploited as templates for the design and development of small peptide-based fungicides with diverse MoA for future use in agriculture (Shafee & Anderson, 2019, Parisi et al., 2019).

## EXPERIMENTAL PROCEDURES

### Fungal cultures and spore suspensions

The fungal strain *Botrytis cinerea* T4 was cultured in V8 (20 %) medium for 7-25 days at 25°C. Fungal spores were harvested by flooding the fungal growth plates with sterile water. The spore suspension was filtered through two layers of Miracloth, centrifuged at 13,600 rpm for 1 min, washed, and re-suspended in low-salt Synthetic Fungal Medium (SFM) (Liang *et al*., 2001). The spore suspension of each fungal pathogen was adjusted to the desired spore density of ∼10^5^ spores/mL using a hemocytometer.

### Peptide synthesis and purification

The peptides GMA4CG_WT and GMA4CG_V1-V6 were chemically synthesized with an initial purity of ∼80-85% from multiple commercial vendors and further purified using a C-18 reverse phase HPLC as described (Velivelli *et al*., 2020b). The HPLC fractions containing the peptide were lyophilized and resuspended in the nuclease-free water. Peptide concentrations were determined using the BCA assay. For peptides containing Trp, peptide concentration was also determined using NanoDrop 2000c Spectrophotometry (Thermo-Fisher Scientific, MA) to confirm concentrations determined by the BCA assay.

### Antifungal activity assays

The minimal inhibitory concentration (MIC) of each peptide was determined using the resazurin cell viability assay (Chadha & Kale, 2015, Li et al., 2019). Briefly, this involved incubation of the pathogen/peptide mixture for 48 h, adding 0.01% (w/v) resazurin solution to each well, and re-incubating overnight. A change in color of the resazurin dye from blue to pink or colorless indicated the presence of live fungal cells. MIC was determined as the minimal concentration of peptide at which no change in color was observed. The MIC value of each peptide was also determined in SFM supplemented with 100 mM NaCl, 100 mM KCl, or 2 mM CaCl2.

### Membrane permeabilization assay using SYTOX^TM^ Green (SG) uptake

The fluorescent dye SG (Thermo-Fisher Scientific, MA) was used to identify GMA4CG_V6-induced membrane permeabilization of *B. cinerea* conidia and germlings as described previously (Li et al., 2019). After exposure to 1.5 µM GMA4CG_V6 for 30 min, *B. cinerea* spores and germlings were imaged using fluorescence microscopy on a Leica SP8-X confocal microscope with an excitation and emission wavelength of 488 and 508-544 nm, respectively.

The GMA4CG_V6 induced membrane permeabilization of conidia was also assessed quantitatively in the absence and presence of 100 mM KCl. Fresh conidia at a concentration of 1× 10^5^/mL were incubated with GMA4CG_V6 at 1×, 2×, and 4× MIC for 1 h in the absence and presence of 100 mM KCl for 1 h and 5 h, respectively. Samples were deposited into glass-bottom petri dishes and imaged by confocal microscopy at an excitation wavelength of 488 nm and an emission wavelength ranging from 508 to 544 nm in the presence of 1 µM SG added 15 min prior to imaging. Fresh conidia incubated with SG alone were used as negative controls. A Leica SP8-X confocal microscope was used for all confocal imaging and Leica LASX (version 3.1.5.) software was used to process the images. The percentage of conidia showing membrane permeabilization was determined by counting the number of cells showing fluorescent signal relative to the total number of cells counted.

The effect of MtDef4, GMA4CG_WT, and GMA4CG_V6 on membrane integrity of *B. cinerea* germlings was determined using a SG (final concentration: 1 µM) uptake quantification assay in the presence and absence of 100 mM KCl (Li et al., 2019). In brief, membrane permeabilization in *B. cinerea* germlings treated with different concentrations (0.187, 0.375, 0.75, 1.5, and 3 µM) of MtDef4, GMA4CG_WT, and GMA4CG_V6 was measured at 5 h in the presence and absence of 100 mM KCl. SG uptake was quantified by measuring the fluorescence in a SpectraMax® M3 spectrophotometer (excitation: 488 nm; cutoff: 515 nm; emission: 530 nm).

### Internalization and subcellular localization of GMA4CG_V6 in *B. cinerea* germlings

Time-lapse confocal laser scanning microscopy was used to monitor uptake and subcellular localization of fluorescently labeled peptide as described previously (Velivelli et al., 2020a). Crude TMR fluorophore labeled GMA4CG_V6 (TMR-GMA4CG_V6) was custom synthesized (Agentide Co, Piscataway, NJ) and purified by HPLC. *B. cinerea* germlings were generated by incubating conidia overnight in SFM medium. They were challenged with TMR-GMA4CG_V6 at concentrations of 0.5, 1.5, or 20 µM and imaged by time-lapse confocal microscopy. The excitation and emission wavelengths for imaging TMR-GMA4CG_V6 were 562 and 580 to 680 nm, respectively.

The sub-localization of GMA4CG_V6 was determined by visualizing its localization with the vacuole-specific dye CMAC and the cell membrane specific dye FM4-64. The excitation and emission wavelengths for CMAC were 353 nm and 405 to 480 nm, respectively, and for FM4-64 were 562 and 680 to 800 nm, respectively. Co-localized pixels (white overlay) of TMR-GMA4CG_V6 and CMAC/FM4-64 was displayed using Leica LAS-X software. The threshold for each channel was 50% and the pixels out of this range were considered uncorrelated. The threshold for the background was set as 20%. Images were captured 30 min after incubation with peptide.

### Quantification of cell wall periphery binding intensity

The spores of *B. cinerea* were treated with 1.5 µM DyLight550-labeled MtDef4, TMR-GMA4CG_WT, and TMR-GMA4CG_V6 for 20 min. Spores of each treatment were imaged using a C-Apochromat 63× water immersion objective lens in apotome mode with a ZEISS Elyra7 super-resolution microscope at excitation and emission wavelengths of 561 and 590 nm, respectively. Images were processed using the SIM module and 3D leap processing with auto sharpness setting in ZEN Black 3.0 SR FP2 software. For each captured cell, the cell surface binding intensity was quantified by free hand drawing a line intensity profile along the cell surface. The fluorescence intensity of each cell wall periphery was normalized by its perimeter length as described (Velivelli et al., 2020a). The lower threshold was set to 7000 to subtract the signal in the cytoplasm.

### Semi-*in planta* antifungal activity assays

The semi-*in planta* antifungal activity of each peptide against *B. cinerea* was determined using *N. benthamiana* Nb1 detached leaves as described previously (Li et al., 2019). Each peptide was tested at concentrations ranging from 0.75, 1.5, 3, and 6 µM. Following incubation of each peptide/fungal spore mixture at room temperature for 48 h, leaves were photographed in white light. High-resolution fluorescence images were also taken using CropReporter (PhenoVation, Wageningen, Netherlands). These images depicted the calculated FV/FM (maximum quantum yield of photosystem II) values of diseased area affected by *B. cinerea* infection. Colors in the images show five different classes ranging from class I to class V (0.000 to 0.700) depicting varying degrees of tissue damage. On one end of the scale, green represents class V (0.600 to ≥0.700) high photosynthesis efficiency. On the other end of the scale, red represents class I (0.000 to 0.160) low photosynthesis efficiency.

### *In planta* antifungal activity assays

To test preventative and curative antifungal activity of the peptides against *B. cinerea*, four-week-old *N. benthamiana* Nb1 plants grown under controlled growth conditions were used. For preventative activity, GMA4CG_WT and GMA4CG_V6 at 12 or 24 µM concentration were sprayed (2 mL/plant) on experimental plants. Control plants were sprayed with 2 mL of water/plant. The peptide solution or water was allowed to dry out on the surface of the leaves. *B. cinerea* spores (∼8×10^4^) suspended in 0.5 × SFM were sprayed on the experimental and control tomato plants (1 mL/plant), transferred to Ziploc plastic boxes under high humidity, and disease symptoms recorded after 48 h. Photographs were taken at 72 h post-inoculation using both white light and CropReporter (PhenoVation, Wageningen, Netherlands). Quantification of symptoms was made by determining maximum quantum yield of PSII (Fv/Fm) as described above for the semi-*in planta* assays.

For the curative antifungal activity tests, *N. benthamiana* Nb1 plants of similar age were first sprayed with *B. cinerea* spores (∼8×10^4^) suspended in 0.5× SFM (1 mL/plant). After 2, 15, and 30 h of pathogen challenge, 2 mL of GMA4CG_WT or GMA4CG_V6 was sprayed on the plants at a concentration of 12 µM. Control plants were each sprayed with 2 mL of water. Disease symptoms were observed after 48 h of pathogen challenge. Plants were photographed at 72 h post infection in white light and quantification was done as described above. The curative antifungal activity of GMA4CG_WT and GMA4CG_V6 at concentration of 6, 12, and 24 µM was tested on 3-week-old tomato (*Solanum lycopersicum* L. cv. Mountain Spring) plants pre-inoculated with *B. cinerea* for 12 h. The severity of disease symptoms was rated using the scoring method for each inoculated plant as follows: 0, no symptoms; 1= 1-25%, 2= 26-50%, 3= 51-75% and 4= 76-100% of leaf surface infected (Segarra *et al*., 2017).

### Visualization of fungal growth on tomato leaf surface

Fresh *B. cinerea* conidia were collected and sprayed on leaves of 3-week-old tomato plants and allowed to germinate for 12 h. After that, GMA4CG_V6 at 6 µM was applied on the same leaves and incubated in a container under high humidity. Leaves applied with spores and sterile water were set as negative controls. After 48 h, leaves were fixed in 4% paraformaldehyde, stained with CFW for 30 min, and images were captured by an Axio Zoom.V16 microscope (ZEISS) with a PlanNeoFluar Z 1.0x objective under tile mode. The excitation wavelength was 335-383 nm, and the emission wavelength was 420-470 nm. Images were visualized and processed by Zen Black 3.0 SR FP2 software (ZEISS).

### Cell death assay of GMA4CG_V6 with propidium iodide

To determine fungicidal activity, freshly harvested conidia of *B. cinerea* were incubated with 1.5 µM GMA4CG_V6 for 1, 6, and 24 h. After incubation, conidia were centrifuged and washed twice in 1x SFM and re-suspended in the same medium. The spores were subsequently allowed to germinate overnight. Microscope images were taken, and germination rate was calculated. Propidium iodide (PI) at 0.5 µg/mL was used as a marker to count dead spores. Conidia of *B. cinerea* were incubated with GMA4CG_V6 for 0.25, 0.5, 1, and 2h. PI was added to the spores/peptide mixture 10 min in advance of detection. Images were captured on the Leica SP8-X confocal microscope (excitation: 535 nm, emission: 600-650 nm).

### Circular dichroism spectroscopy

A steady-state wavelength circular dichroism spectrum was collected for GMA4CG_WT and GMA4CG_V6 (∼0.05 mM peptide, 20 mM sodium acetate, pH 5.6) on a calibrated Aviv Model 410 spectropolarimeter (Lakewood, NJ). Using a quartz cell of 0.1 cm path length and a temperature of 20 °C, the data was recorded in 0.5 nm increments between 200 and 260 nm with a bandwidth of 1.0 nm and a time constant of 1.0 s. The reported steady-state wavelength spectrum was the result of averaging two consecutive scans, subtracting a blank spectrum, and automatic line smoothing using Aviv software.

### Antifungal activity of the alanine scanning variants of GMA4CG_V6 with no disulfide bond

The minimal inhibitory concentration (MIC) of linear GMA4CG_V6 alanine scanning variants was determined using the resazurin cell viability assay as described in the methods section. The MIC value of all variants was also determined in SFM supplemented with 100 mM NaCl.

### Cell wall/surface localization analysis of GMA4CG_V6

*B. cinerea* germlings were treated with 1.5 µM of TMR-GMA4CG_V6 and, after 5 min, CFW (5 µM) was added into the same petri dish. Samples were observed using a confocal (Leica SP8-X) and super-resolution microscope (Zeiss Elyra7). To avoid bleed-through of the two fluorescent signals, confocal images were captured sequentially using a fast line switch configuration (TMR-GMA4CG_V6 excitation: 542 nm, emission: 580-680 nm, and CFW excitation: 405 nm, emission: 410-557 nm). To obtain higher resolution images, the C-Apochromat 63×1.4 oil immersion objective lens in apotome mode was used with the ZEISS Elyra7 super-resolution microscope (Zeiss). The excitation wavelength was 405 nm for the CFW signal and 561 nm for the TMR signal with emission wavelengths of 420-480 and 570-620 nm, respectively. Images were processed using the SIM module plus 3D and leap processing with autosharpness. The channels were aligned based on 0.1 µm TetraSpeck™ microspheres (T7279, Thermo Fisher Scientific) using ZEN Black 3.0 SR FP2 software.

### Fluorescence Redistribution After Photobleaching (FRAP) experimental setup and analysis

The FRAP experiments were performed on a Leica SP8-X microscope (FRAP Wizard Module, LAS X Version 3.1.5.16.16308) using the HC PlanApochromat 63× water immersion objective len (Numerical aperture = 1.2) with the pinhole set to 1 Airy unit and zoom set to 1.77 (Velivelli et al., 2020a). Images were acquired for the selected cellular regions (germling cytoplasm, vacuole, cell periphery, and spore cell wall/periphery) after 1 h incubation with GMA4CG_V6 under the following identical imaging and FRAP conditions: 10 pre-bleach and 700 post-bleach frames (512 x 64 pixels, pixel size = 204 nm) captured at 173 ms intervals without averaging using 550 excitation and 560-625 nm emission wavelengths with PMT detectors and a transmitted light channel. The power of the 550 nm laser was set at 3.5% during imaging to prevent detectable photobleaching for the duration of each FRAP experiment (∼3 min). A 1.52 µm^2^ circular bleach region was applied with 100% laser excitation power (550 nm wavelength) for 3.5 s immediately following pre-bleach and prior to post-bleach imaging. FRAP analysis was performed on 24 data sets for each subcellular region showing no evidence of sample drift during acquisition. A single exponential curve fitting function was applied post-acquisition to extract the half-maximum tau *(τ½),* mobile fraction, and immobile fraction for each condition.

### Phospholipid binding assays

His-tagged GMA4CG_WT and GMA4CG_V6 was obtained from Biomatik Inc. (Ontario, Canada) and further purified by HPLC. The His-tagged peptides (GMA4CG_WT and GMA4CG_V6) phospholipid binding assays were performed using the PIP Strips^TM^ (Echelon Biosciences Inc., CA) as previously described (Islam *et al*., 2017). Hexa-His peptide (Genscript, Inc., NJ) was used as a control. The peptide binding to the membrane strips was detected using 0.5 µg/mL His-tag monoclonal antibody purchased from Thermo-Fisher Scientific, MA. The binding signals were detected using the SuperSignal West Pico Chemiluminescent Substrate Kit (Thermo-Fisher Scientific, MA) following the manufacturer’s protocol. Quantitative analysis of the signal was performed using Image J (https://imagej.nih.gov/ij/download.html).

### The binding of GMA4CG_WT and GMA4CG_V6 on tomato leaf surfaces

The fluorophore-labeled peptides, TMR-GMA4CG_WT and TMR-GMA4CG_V6, were applied on the surface of tomato leaves and allowed to air dry for 1 h. The leaf samples were subjected to microscopic observation using a Leica SP8-X confocal microscope (TMR wavelength: excitation: 562 nm; emission: 580-680 nm, and autofluorescence of tomato cells wavelength: excitation: 405 nm; emission: 416-483 nm) with an APO CS2 40× water objective lens before and after a gentle rinse twice in sterile water. The average signal intensity from 10-12 ROIs of each image was collected for quantification and 26-32 images were captured. The signal intensity of the after-wash-sample was normalized by the signal of the sample before washing.

### Statistical analysis

All experiments were performed in three or more technical replicates where appropriate and in three independent experiments. The phospholipid binding assays and quantitative measurements of cell membrane permeability in *B. cinerea* were performed using two independent biological replications. Data were graphed and analyzed in GraphPad Prism (version 8.0 or 9.4.1) software using unpaired, two-tailed Student’s *t* tests, and one-way ANOVA with Tukey’s honestly significant difference test (HSD) for comparison of multiple groups. Statistical significance was accepted at *p* < 0.05.

## Acknowledgements

This work was supported, in part, by the National Science Foundation-EAGER grant IOS:1955461 awarded to D.S. We acknowledge imaging support from the Advanced Bioimaging Laboratory (RRID:SCR_018951) at the Danforth Plant Science Center and usage of the Leica SP8-X confocal microscope acquired through an NSF Major Research Instrumentation grant (DBI-1337680) and usage of the ZEISS Elyra 7 Super-Resolution Microscope acquired through an NSF Major Research Instrumentation grant (DBI-2018962). We are thankful to Dr. Saurav Misra of the Danforth Center for his attempts to generate structures of the peptides using AlphaFold. This work was funded by the Wells Fargo IN^2^ grant awarded to Peptyde Bio. Part of this research was performed at the Pacific Northwest National Laboratory (PNNL), a facility operated by Battelle for the U.S. Department of Energy, including access to the W.R. Wiley Environmental Molecular Sciences Laboratory (EMSL), a national scientific user facility sponsored by the U.S. DOE Biological and Environmental Research program. Battelle operates PNNL for the U.S. Department of Energy under contract DE-AC05-76RL01830.

## Author Contributions

Conceptualization: Meenakshi Tetorya, Hui Li, Kirk J. Czymmek, Dilip M. Shah Funding acquisition: Dilip M. Shah

Methodology: Meenakshi Tetorya, Hui Li, Kirk J. Czymmek, Arnaud Thierry Djami-Tchatchou, Garry W. Buchko, Dilip M. Shah

Project administration: Dilip M. Shah Supervision: Kirk J. Czymmek, Dilip Shah Validation: Kirk J. Czymmek, Dilip Shah

Writing-original draft: Meenakshi Tetorya, Hui Li, Garry Buchko

Writing-review & editing: Dilip M. Shah, Kirk J. Czymmek, Garry W. Buchko

## Supporting information

**S1_Fig.**
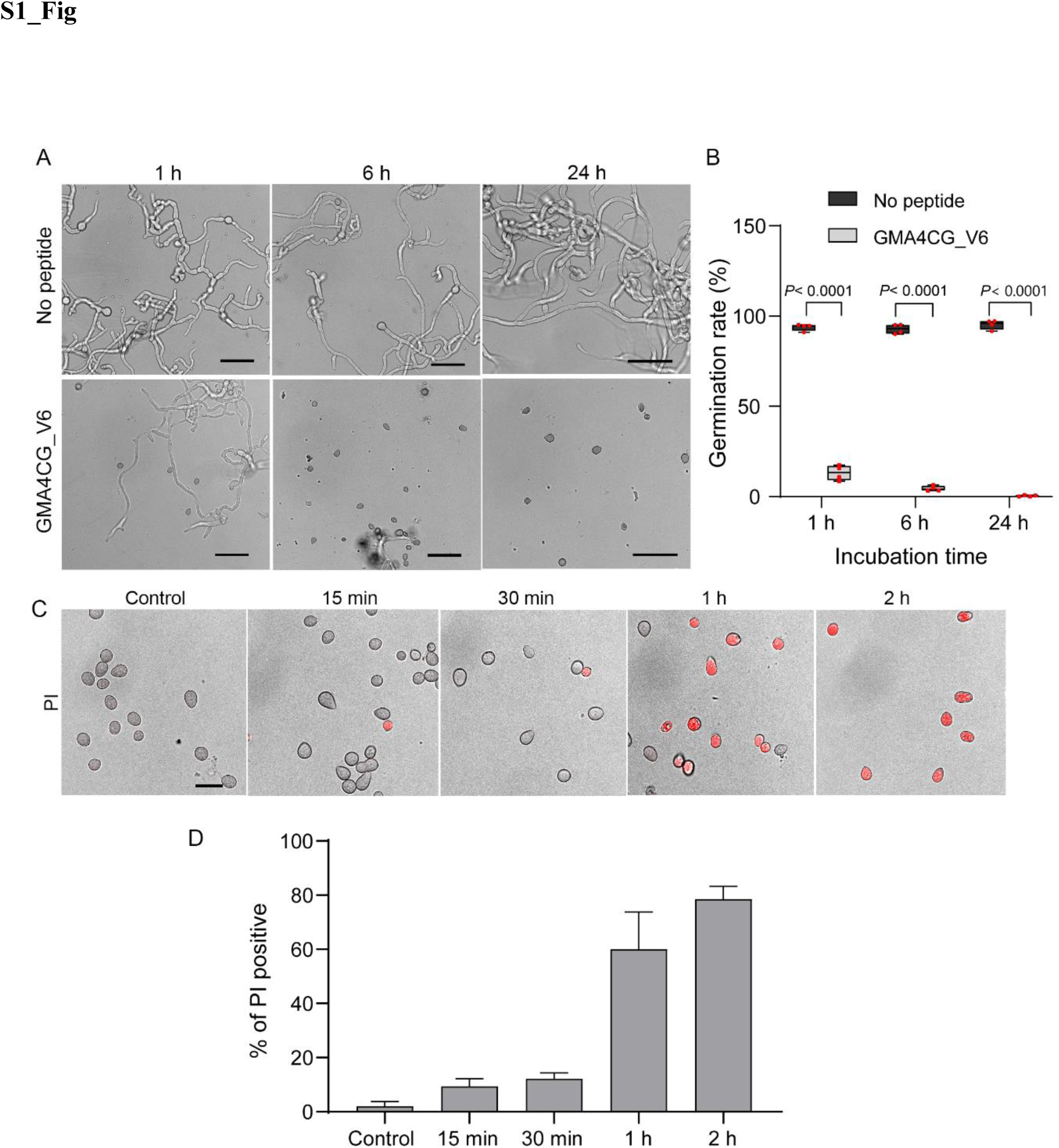
GMA4CG_V6 exhibits fungicidal activity. (**A**) Transmitted light images of the germination and growth of *B. cinerea* spores treated with 1.5 µM GMA4CG_V6 for 1, 6, and 24 h. After each treatment, unbound peptide was removed, and the spores allowed to germinate overnight. Scale bar = 50 µm. (**B**) Quantification of the germination rate of GMA4CG_V6 treated *B. cinerea* spores. Spores treated with sterile water served as controls. Data were collected from four independent replications (n = 4) and plots are shown in box plot. Individual data were overlaid on plots as red points. Statistical analysis was performed by two-sided Student’s t test. (**C**) *B. cinerea* spores were treated with 1.5 µM GMA4CG_V6 and the dead cell staining dye PI for various times. The overlaid images are shown for 0.25, 0.5, 1, and 2 h treatments. Scale bar = 20 µm. (**D**) Quantitative analysis of the percentage of spores stained with PI dye. The top of the columns represents the mean, and the horizontal bars represent the standard deviation (n = 3).

**S2_Fig.**
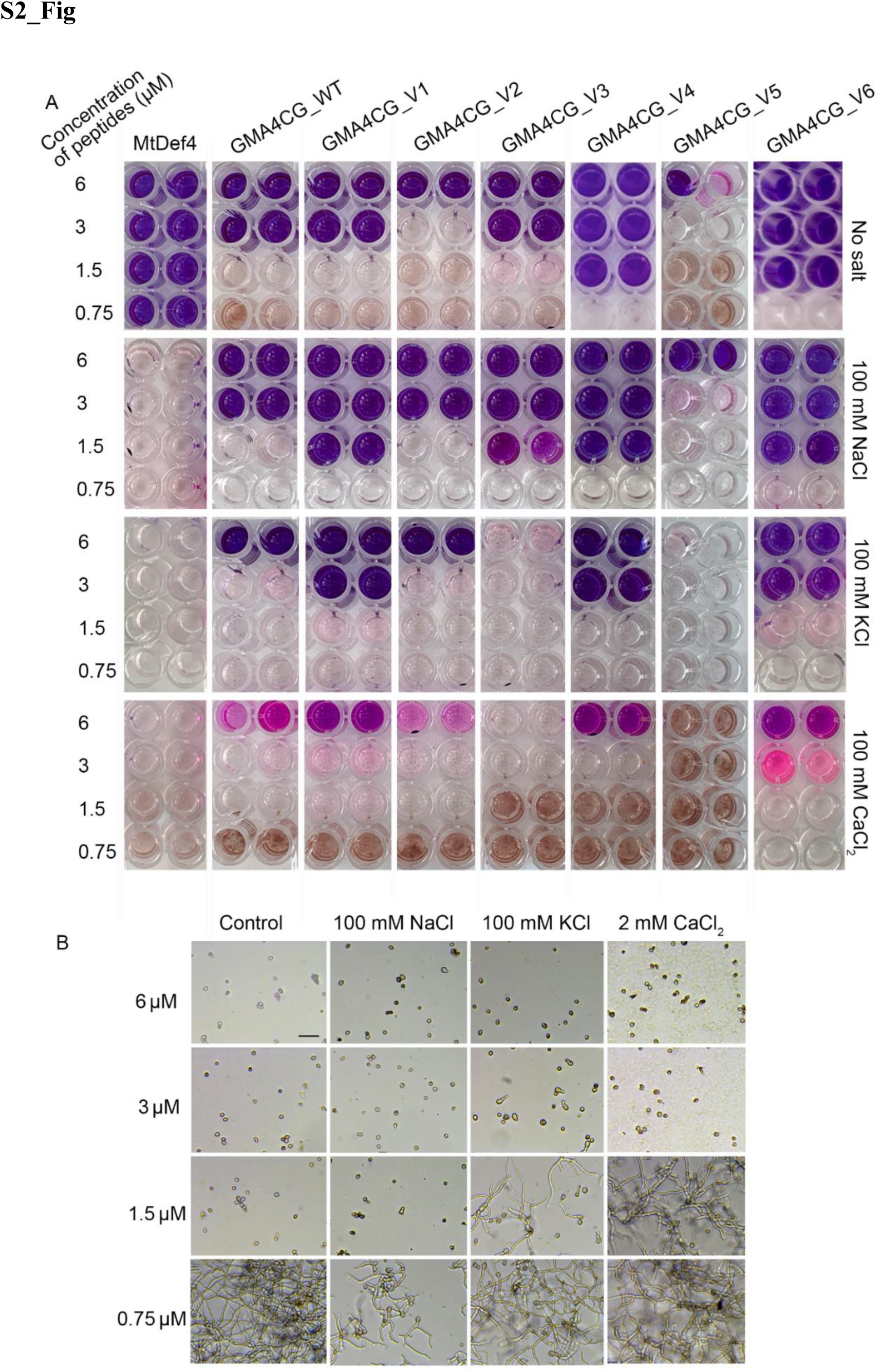
GMA4CG_V6 retains antifungal activity in the presence of salts. **(A)** Antifungal activity of the GMA4CG variants and MtDef4 against *B. cinerea* was determined in 1 x SFM in the absence or presence of 100 mM NaCl, 100 mM NaCl, or 2 mM CaCl2. After 48 h incubation, resazurin dye was added to each well to determine cell viability/cell death. A change from blue to pink/colorless indicated viable fungal cells. **(B)** Microscopic images of *B. cinerea* spores challenged with various concentrations of GMA4CG_V6 in the presence of different salts (Scale bar = 50 μm).

**S3_Fig.**
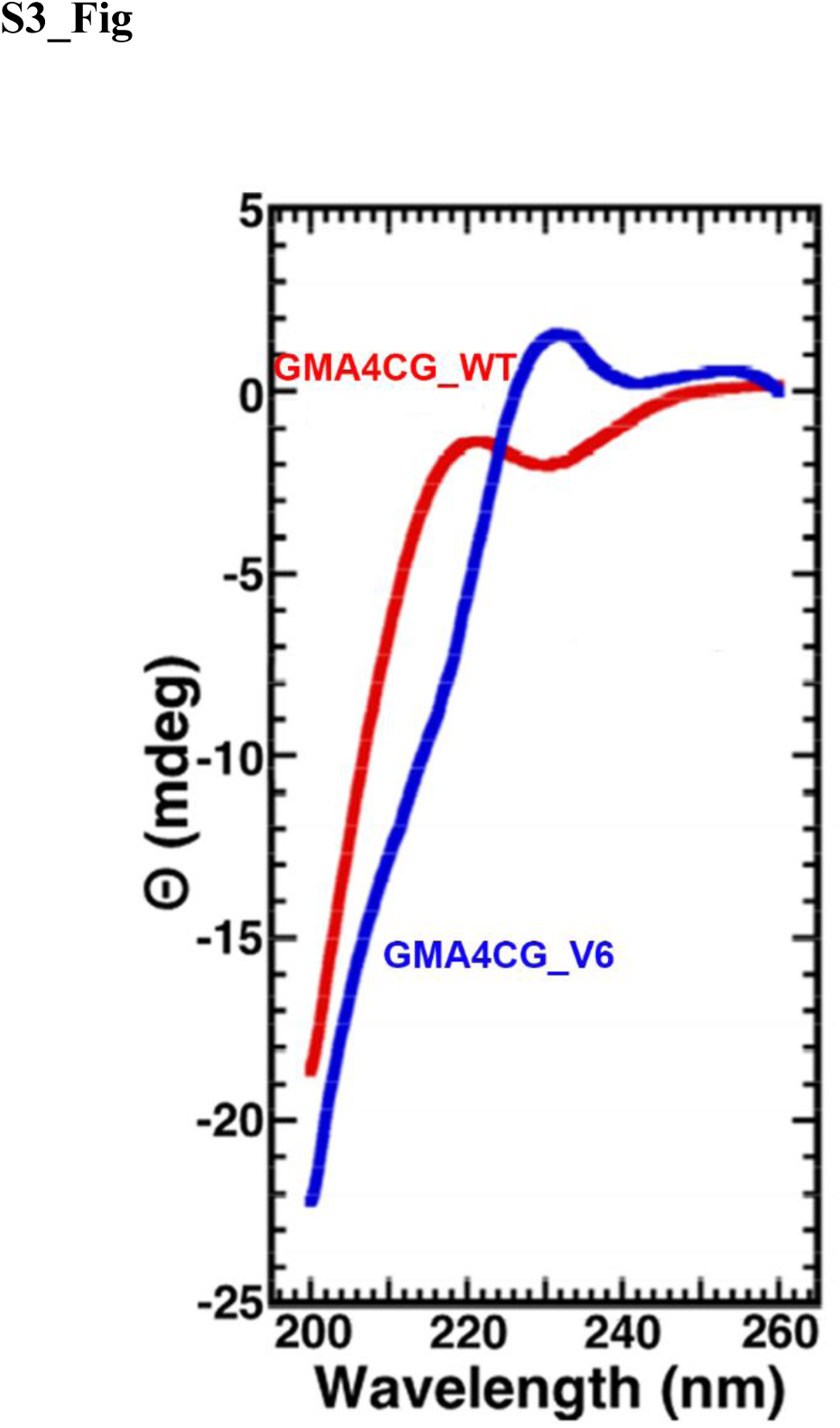
Steady-state wavelength CD spectra for GMA4CG_WT (red) and GMA4CG_V6 (blue) in 20 mM sodium acetate, pH 5.6, at 20 °C.

**S4_Fig.**
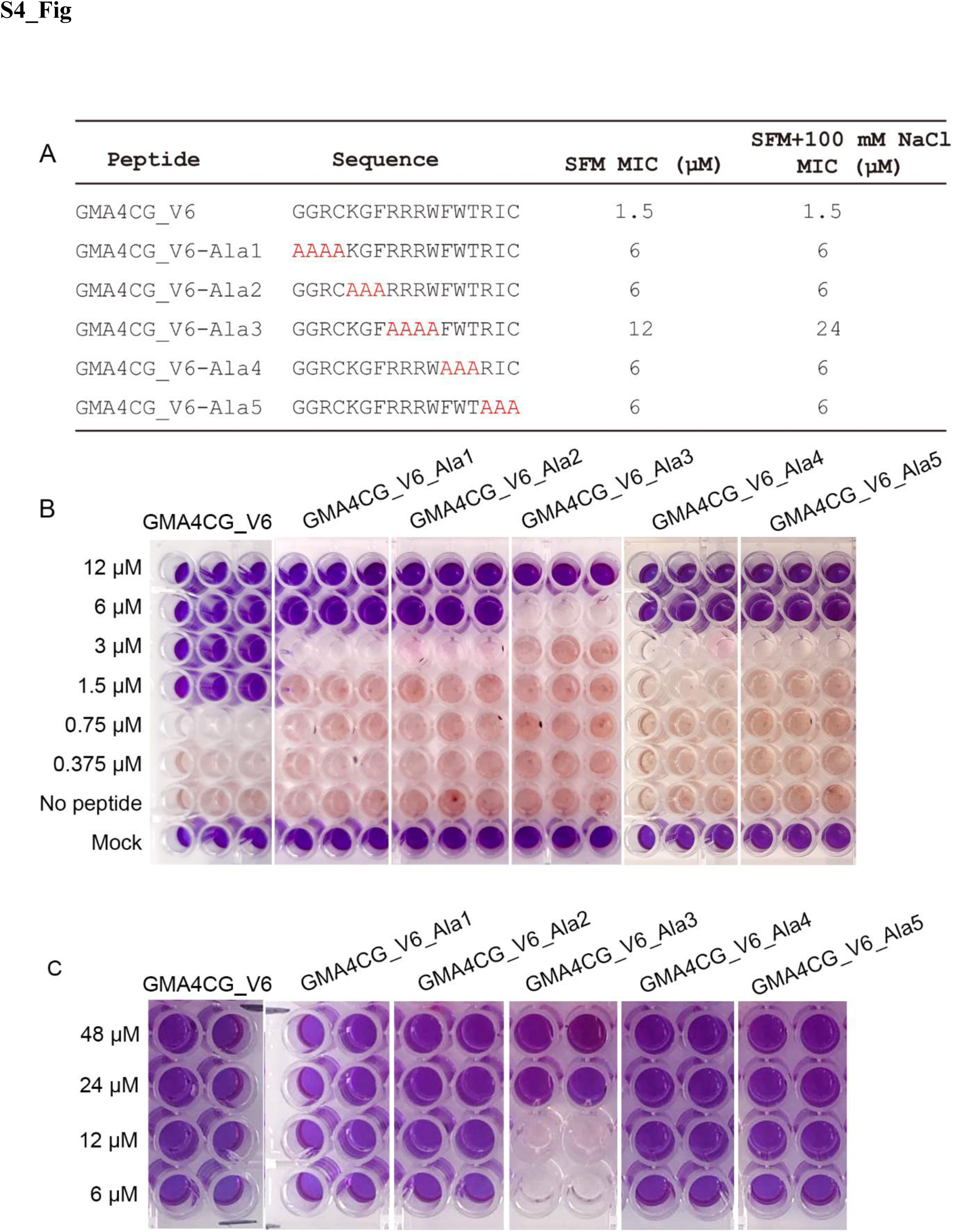
Antifungal activity of the alanine scanning variants of linear GMA4CG_V6 as determined by the resazurin cell viability/cell death assay. (**A**) The primary amino acid sequences for GMA4CG_V6 and the five Ala scanning mutants and their associated MIC values in the absence and presence of salt. The GMA4CG_V6 substitutions are highlighted red. (**B**) Resazurin dye was used to determine cell viability/cell death. A change from blue to pink/colorless indicated viable fungal cells. Antifungal activity of each variant was performed in 1 x SFM. There was an eight-fold loss in activity with GMA4CG_V6_Ala3 whereas all other variants showed a four-fold loss. (**C**) Resazurin cell viability/cell death assay showing the MIC values of alanine scanning variants of GMA4CG_V6 in the presence of 100 mM NaCl. MIC values of all other variants remained unchanged except for GMA4CG_V6_Ala3, which increased another two-fold.

**S5_Fig.**
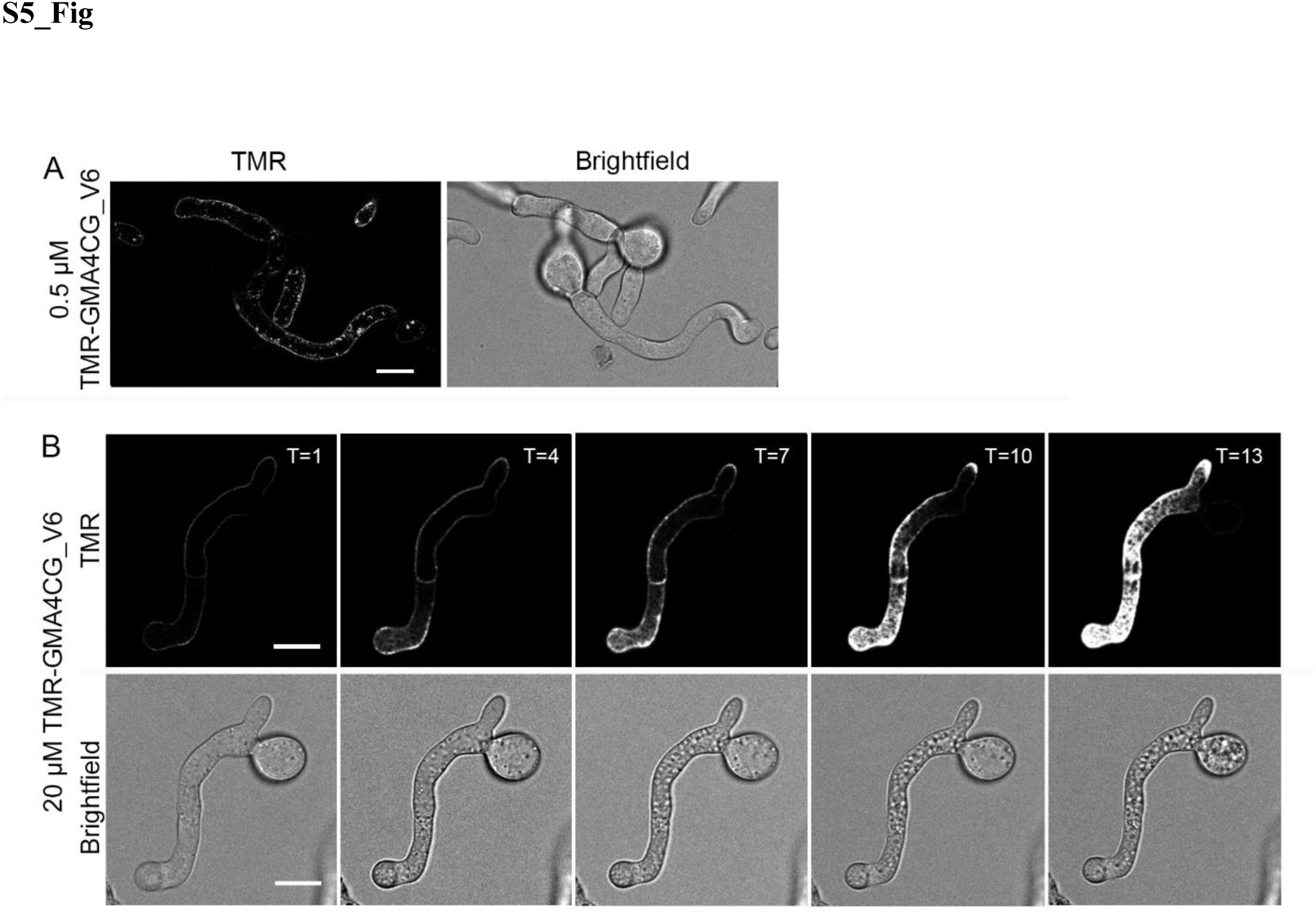
Internalization of TMR-GMA4CG_V6 into *B. cinerea* germlings at sublethal or very high concentrations. **(A)** Confocal microscopy images show internalization of 0.5 µM TMR-GMA4CG_V6. Images were captured 60 min after peptide challenge. **(B)** Time-lapse confocal microscopy images of *B. cinerea* show internalization of 20 µM TMR-GMA4CG_V6 over time (T in min) with images captured every 3 min shown. Scale bars = 10 µm.

**S6_Fig.**
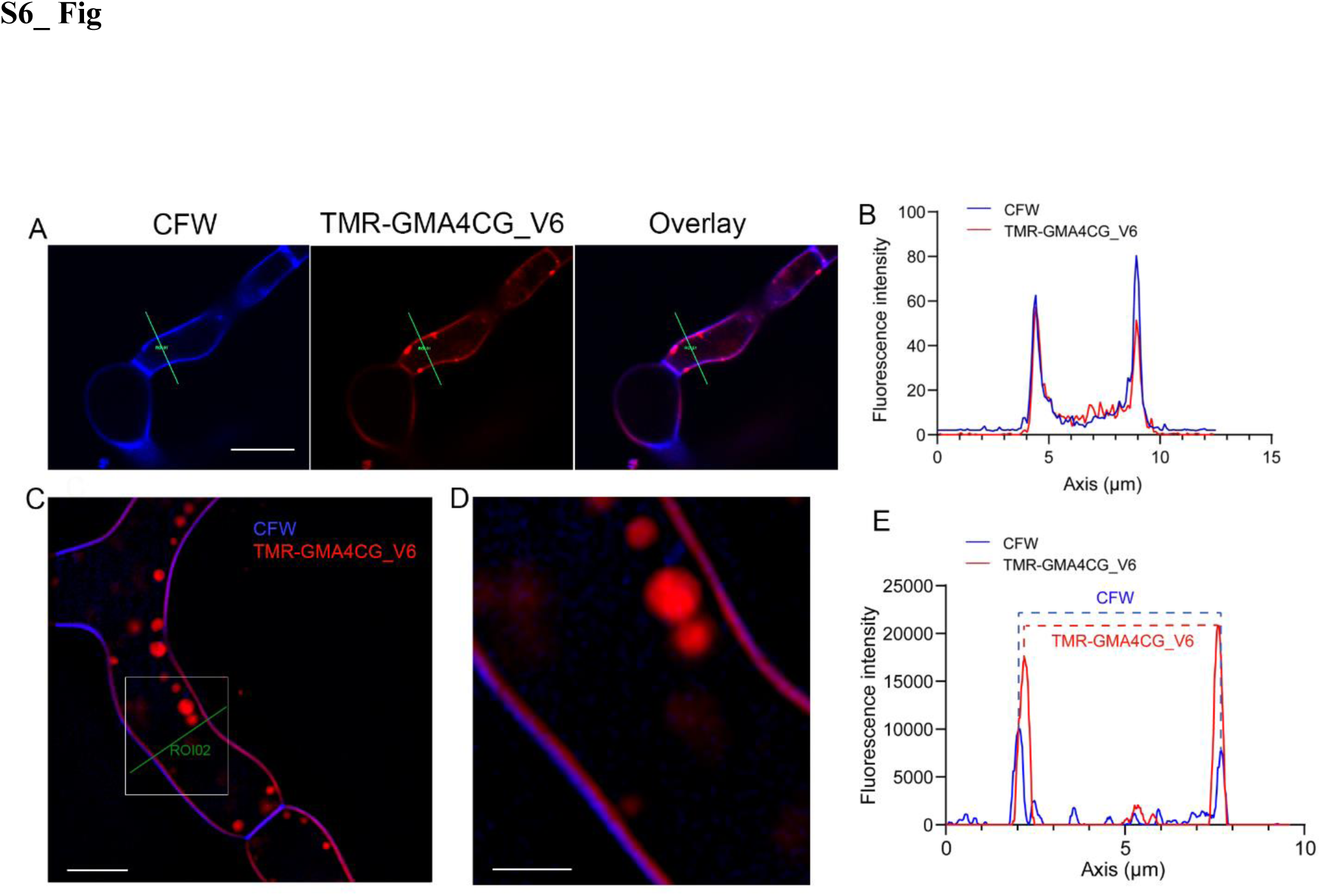
Co-localization of TMR-GMA4CG_V6 and CFW. (**A & C)** Representative fluorescent (CFW and TMR) and channel merged images of *B. cinerea* germlings treated with TMR-GMA4CG_V6 and captured by a confocal microscope (A) and the overlay image by lattice SIM^2^ super resolution microscope (C). Scale bar is 10 µm in A and 5 µm in C. **(B & E)** The fluorescent signal intensity of the ROIs in **A** and **C**, respectively. The dash lines in **E** indicate the shift between the peak of the CFW and TMR signals. **(D)** Enlarged image from the framed area in C. Scale bar = 2 µm.

**S7_Fig.**
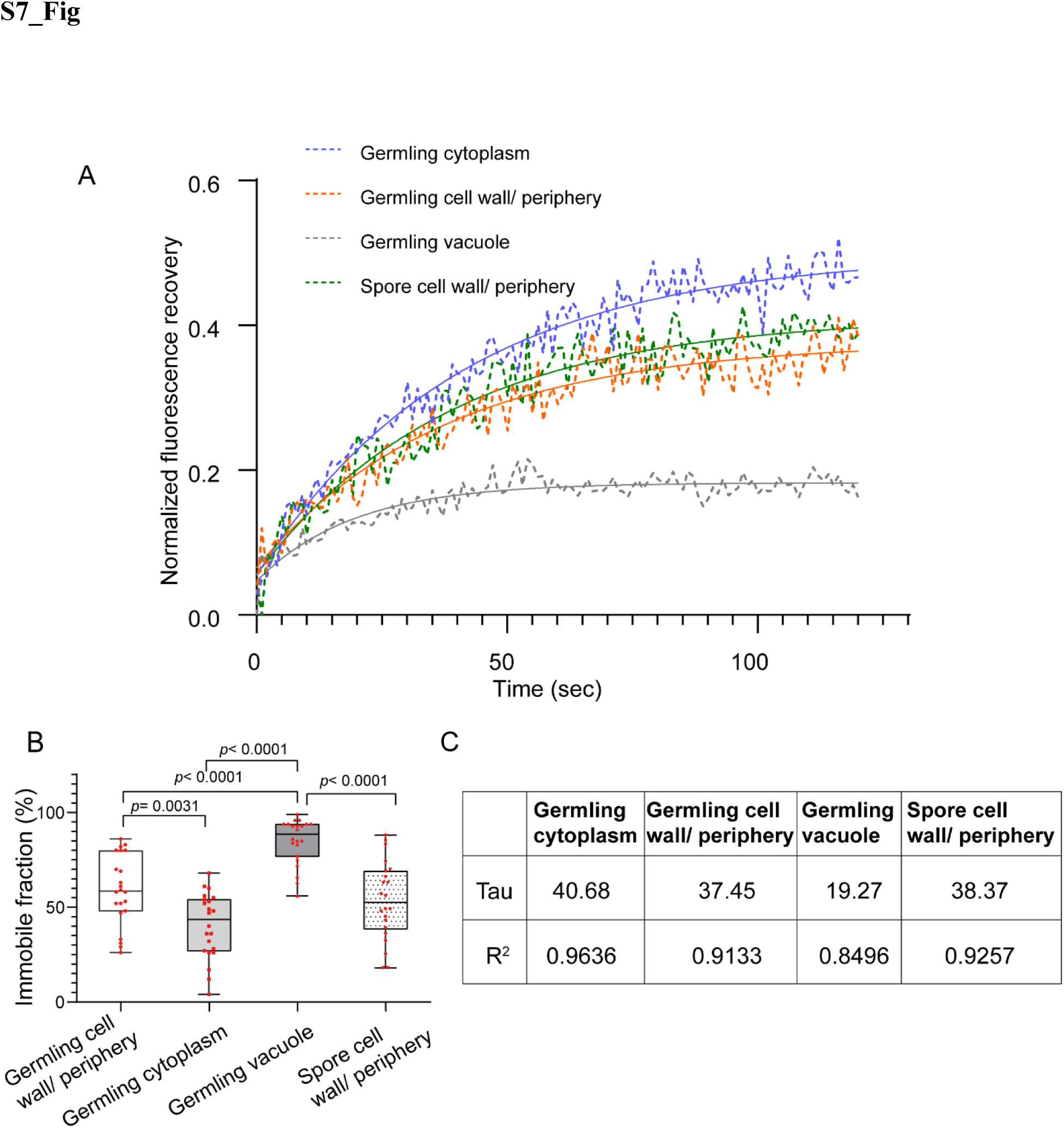
FRAP analysis of the subcellular localization of GMA4CG_V6 in *B. cinerea* germlings and conidia. **(A)** FRAP plots show the normalized fluorescence recovery of TMR-GMA4CG_V6 in germlings cytoplasm (blue), cell walls/ periphery (orange), and vacuoles (gray) and in conidia cell walls/periphery (green) over 120 s. A one-phase association parameter was used to simulate the fitting curve (solid lines) based on the scatter diagram (GraphPad Prism 8.0). Higher recovery rates indicate greater overall mobility. **(B)** The immobile fraction (%) of GMA4CG_V6 localized into the key subcellular compartments of *B. cinerea*. Data are represented using box plots with individual data overlaid as red points. Data analyzed for each subcellular compartment of *B. cinerea* are from n = 20 collections from at least three independent experiments. Asterisks represent significant differences between different subcellular compartments. One-way analysis of variance (ANOVA) was used to determine the statistical difference with Tukey’s honestly significant difference test (HSD) for comparison of multiple groups. **(C)** A table showing the half-time of recovery (Tau) of GMA4CG_V6 localized into the key subcellular compartments of *B. cinerea*. The Tau and R^2^ values for each peptide were calculated using a one-phase association parameter.

**S8_Fig.**
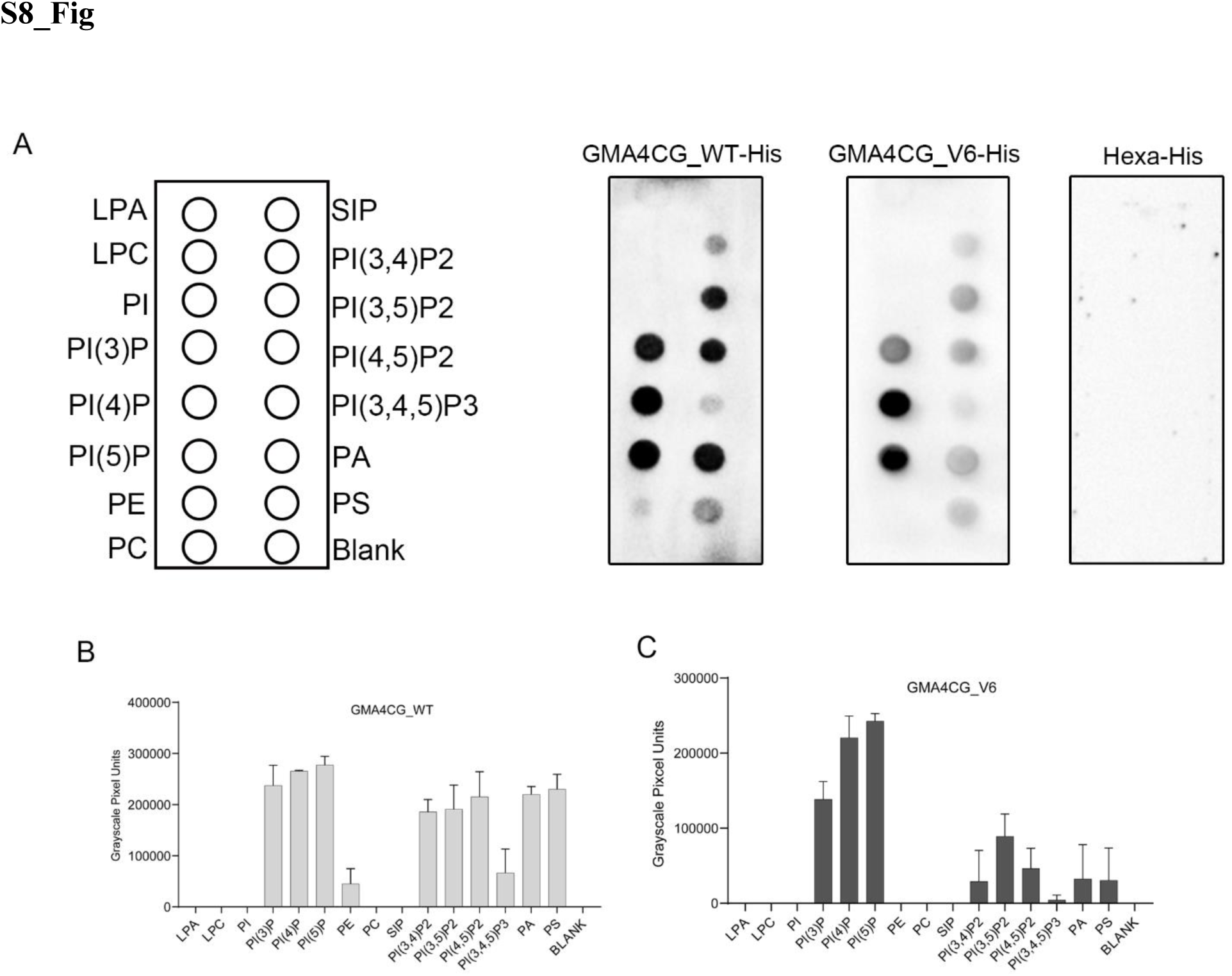
GMA4CG_V6 binding to phospholipids. **(A)** Lipid overlay assays of GMA4CG_WT, GMA4CG_V6, and Hexa-His. Both GMA4CG_WT and GMA4CG_V6 show strong binding to phosphatidylinositol monophosphates PI(3)P, PI(4)P, and PI(5)P while GMA4CG_WT shows strong binding to additional phospholipids including PA and the phosphatidylinositol diphosphates PI(3,5)P2 and PI(4,5)P2. In contrast, GMA4CG_V6 binds to multiple phospholipids, including phosphatidylinositol di/triphosphates PI(3,4)P2, PI(3,4,5)P3, and PA, but rather weakly. **(B & C)** Integrated density (grayscale pixel units) analysis of PIP strip probed with GMA4CG_WT and GMA4CG_V6. Data are means ± SEM of two independent biological replicates (n = 2).

**S9_Fig.**
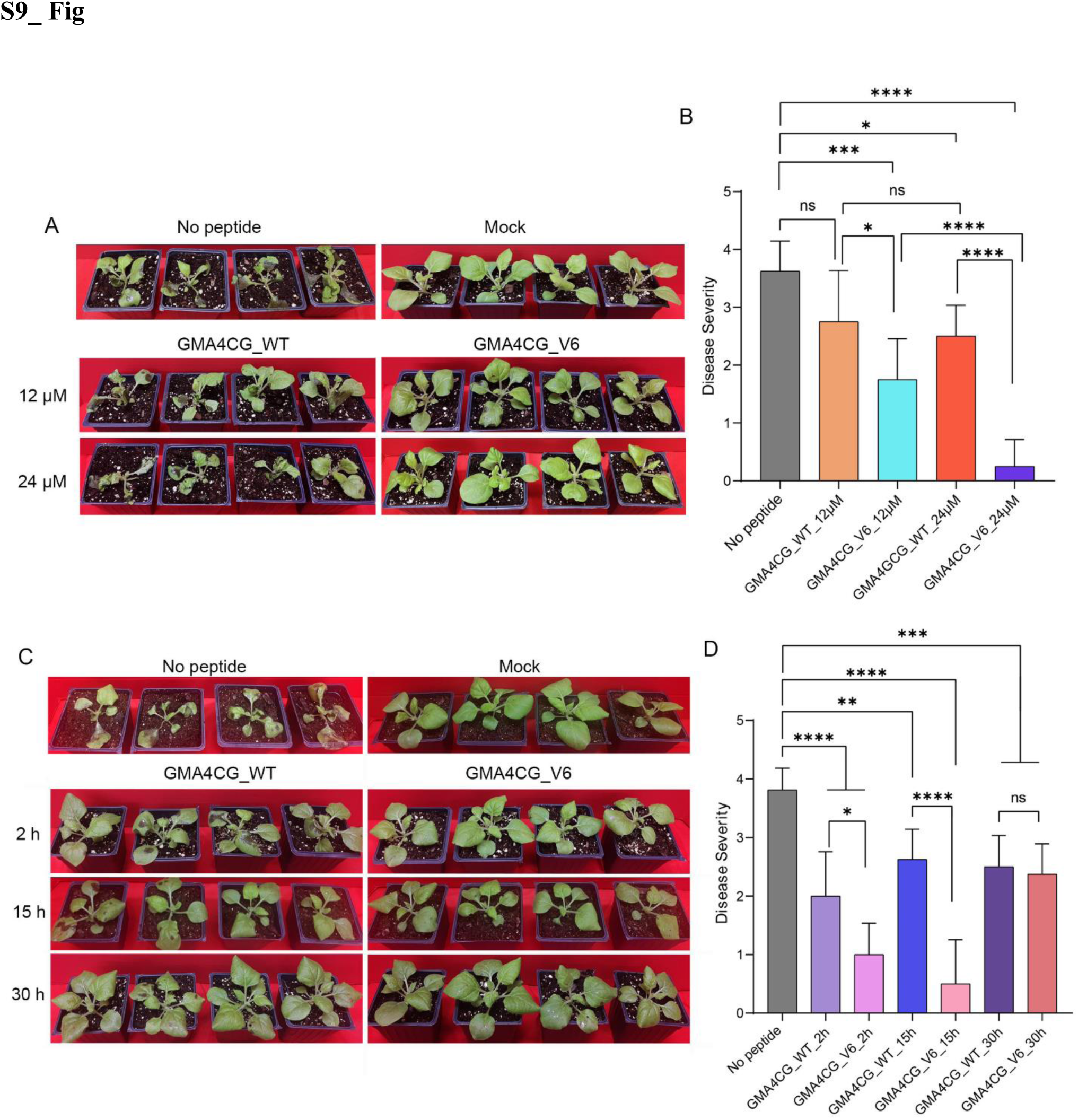
GMA4CG_V6 has preventative and curative antifungal activity against gray mold in *N. benthamiana* plants. **(A)** Preventative activity of GMA4CG_V6. Four-week-old *N. benthamiana* plants were sprayed with either 2 ml water or peptide (GM4CG_WT or GMA4CG_V6) at 12 or 24 µM concentration prior to spray-inoculation with a 1 ml suspension of 8 × 10^4^ *B. cinerea* fungal spores. Plants were observed for symptoms at 72 hpi and photographed in white light. **(B)** Disease severity of tobacco plants inoculated with *B. cinerea* at 72 hpi. The disease severity was rated on a scale of 0 to 4 with 0 = no symptoms; 1 = 1-25%; 2 = 26-50%; 3 = 51–75%; and 4 = 75-100% of the leaf surface infected. One-way analysis of variance (ANOVA) was used to determine statistical differences between control (*B. cinerea*), mock-treated, and peptide-treated plants (GraphPad Prism-version 9.4.1). Four *N. benthamiana* plants (n = 4) for each technical replication were tested in three biological replications. Asterisks represent significant differences between different groups with P values determined (^ns^*p*> 0.05, \**p*≤ 0.05 \*\**p*< 0.01, \*\*\*\**p*≤ 0.0001) using Tukey’s honestly significant difference test for comparison of multiple groups. **(C)** Curative antifungal activity of GMA4CG_V6. Four-week-old *N. benthamiana* plants were spray-inoculated with a 1 mL suspension of 8 × 10^4^ of *B. cinerea* fungal spores at 2, 15, or 30 h and then sprayed with 2 mL of 12 µM GMA4CG_WT, 12 µM GMA4CG_V6, or water. Plants were photographed in white light after 72 h. **(D)** Disease severity was rated as described above. Four *N. benthamiana* plants (n = 4) for each technical replication were tested in three biological replications. Asterisks represent significant differences between different groups with *p* values determined (^ns^*p*> 0.05, \**p*≤ 0.05 \*\**p*< 0.01, \*\*\*\**p*≤ 0.0001) using Tukey’s honestly significant difference test for comparison of multiple groups.

**S10_Fig.**
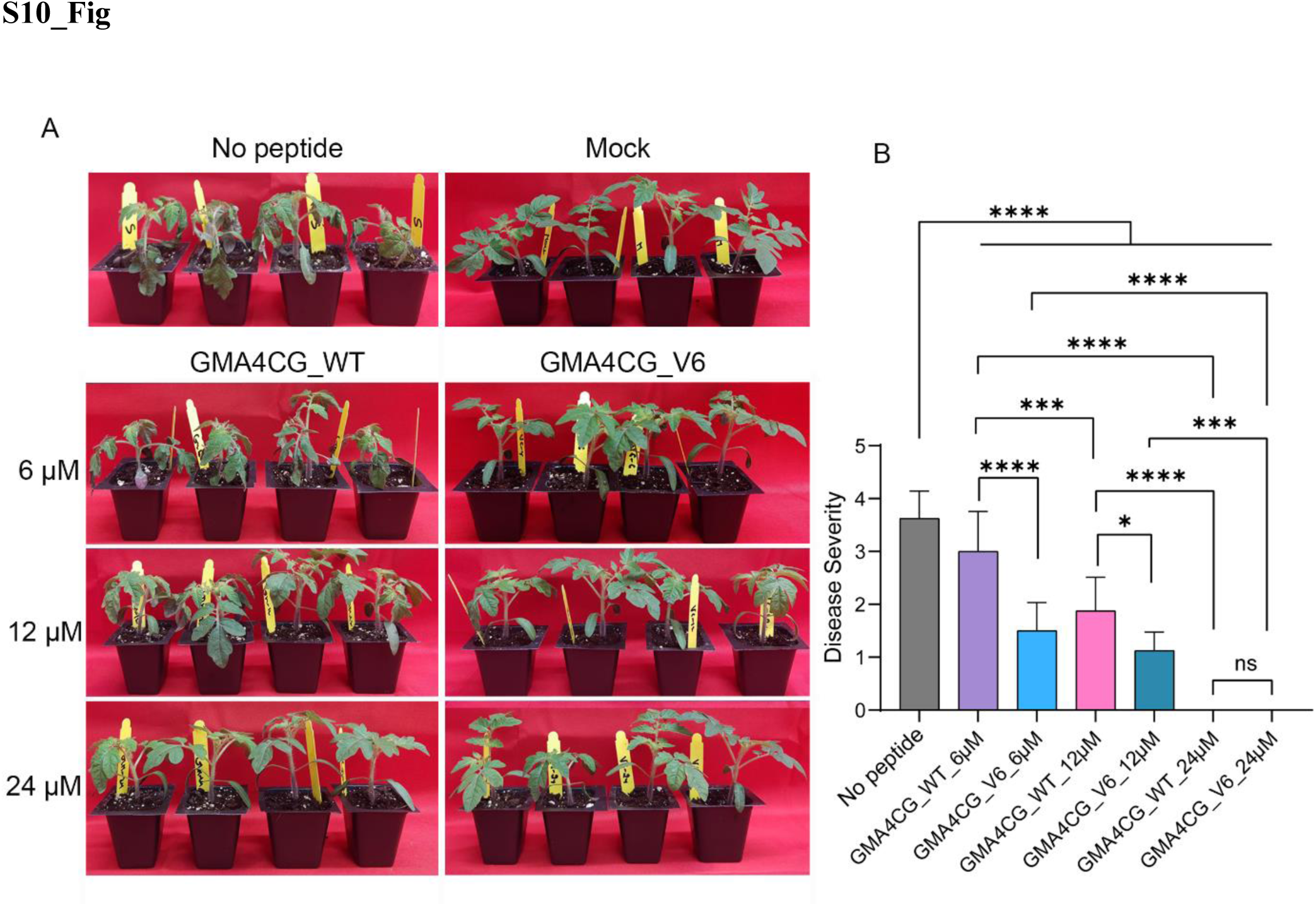
GMA4CG_V6 has curative antifungal activity against gray mold in tomato plants. **(A)** Three-week-old tomato (*Solanum lycopersicum* L cv. Mountain Spring) plants were spray-inoculated with a 1 ml suspension of 8 × 10^4^ fungal spores and subsequently sprayed 12 h later with 2 ml of GMA4CG_WT or GMA4CG_V6 at a concentration of 6, 12, or 24 µM or with 2 ml of water (controls). Plants were scored for gray mold symptoms at 72 hpi and photographed in white light. **(B)** Disease severity of tomato plants inoculated with *B. cinerea* 72 hpi. The disease severity was rated on a scale of 0 to 4 with 0 = no symptoms; 1 = 1-25%; 2 = 26-50%; 3 = 51– 75%; and 4 = 75-100% of the leaf surface infected. One-way analysis of variance (ANOVA) was used to determine statistical differences between control (*B. cinerea*), mock, and peptide experiments (GraphPad Prism-version 9.4.1). Four tomato plants (n = 4) for each technical replication were tested in three biological replications. Asterisks represent significant differences between different groups with *p* values (^ns^*p*> 0.05, \**p*≤ 0.05 \*\**p*< 0.01, \*\*\*\**p*≤ 0.0001) determined using Tukey’s honestly significant difference test for comparison of multiple groups.

**S11_Fig.**
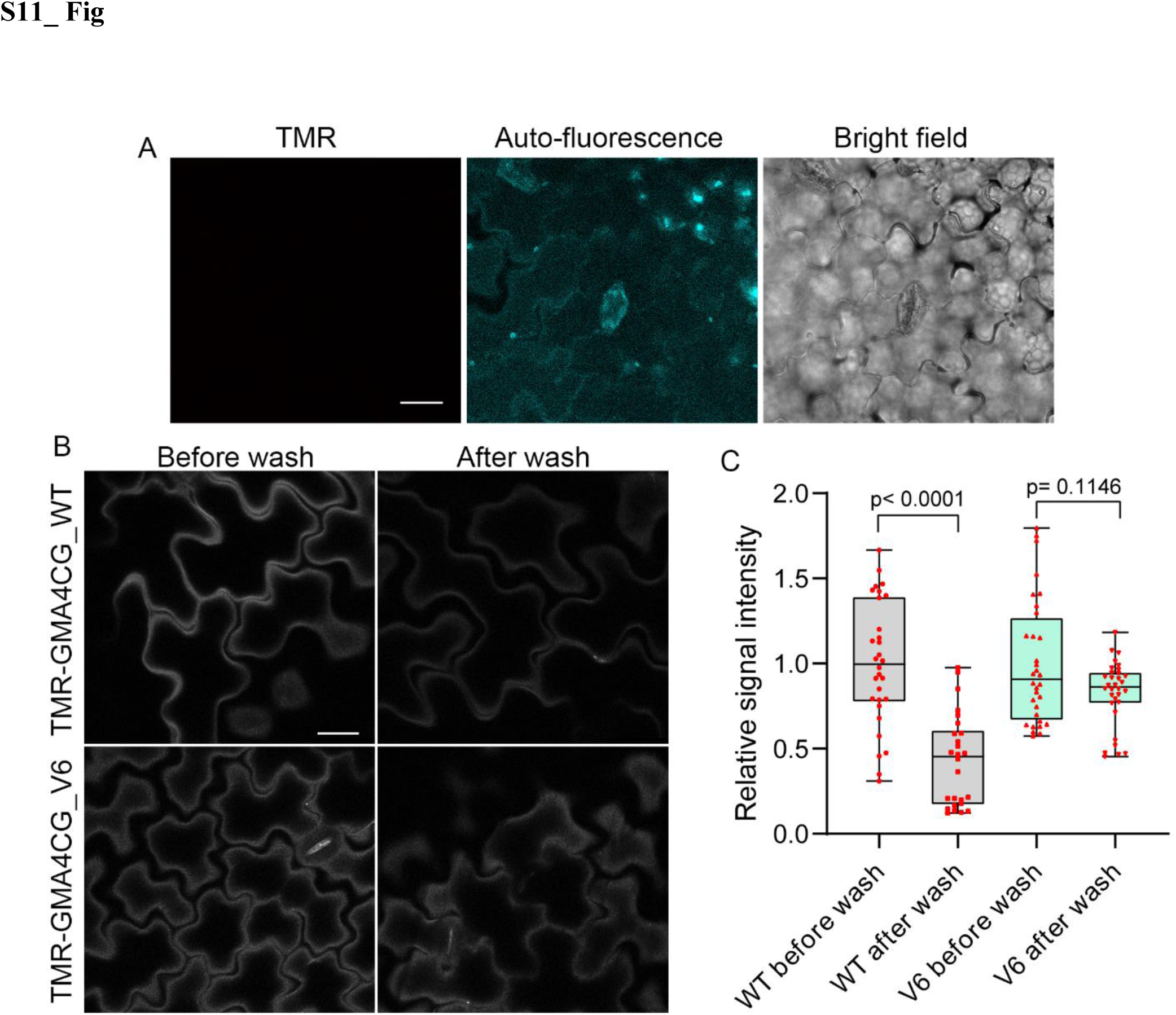
Binding of TMR-GMA4CG_WT and TMR-GMA4CG_V6 to tomato leaf surfaces. **(A)** Representative confocal microscopy images of tomato leaves with no peptide applied showing the absence of TMR fluorescence signal, background cell wall autofluorescence (cyan), and transmitted light. Scale bar = 20 µm. **(B)** Confocal microscopy images of plant cell walls stained with TMR-GMA4CG_WT and TMR-GMA4CG_V6 showing no signal observation in the cytoplasm. Scale bar = 20 µm. **(C)** Comparison of TMR signal intensity at the tomato cell wall before and after washing with sterile water. The average signal from leaves treated with each peptide before washing served as the control (100%). Each dot indicates individual data points. Asterisks represent significant differences between different groups (unpaired *t* test, n = 26-32) (GraphPad Prism 8.0).

